# Phylogenetic reconstruction based on synteny block and gene adjacencies

**DOI:** 10.1101/840942

**Authors:** Guénola Drillon, Raphaël Champeimont, Francesco Oteri, Gilles Fischer, Alessandra Carbone

## Abstract

Gene order can be used as an informative character to reconstruct phylogenetic relationships-between species independently from the local information present in gene/protein sequences.

PhyChro is a reconstruction method based on chromosomal rearrangements, applicable to a wide range of eukaryotic genomes with different gene contents and levels of synteny conservation. For each synteny breakpoint issued from pairwise genome comparisons, the algorithm defines two disjoint sets of genomes, named partial splits, respectively supporting the two block adjacencies defining the breakpoint. Considering all partial splits issued from all pairwise comparisons, a distance between two genomes is computed from the number of partial splits separating them. Tree reconstruction is achieved through a bottom-up approach by iteratively grouping sister genomes minimizing genome distances. PhyChro estimates branch lengths based on the number of synteny breakpoints and provides confidence scores for the branches.

PhyChro performance isevaluatedon two datasets of 13 vertebrates and 21 yeast genomes by using up to 130 000 and 179 000 breakpoints respectively, a scale of genomic markers that has been out of reach until now. PhyChro reconstructs very accurate tree topologies even at known problematic branching positions. Its robustness has been benchmarked for different synteny block reconstruction methods. On simulated data PhyChro reconstructs phylogenies perfectly in almost all cases, and shows the highest accuracy compared to other existing tools. PhyChro is very fast, reconstructing the vertebrate and yeast phylogenies in less than 15 min.

**Availability:** PhyChro will be freely available under the BSD license after publication

## Introduction

Today, phylogenies of many species can be reconstructed using sequences from numerous proteins, but, despite the availability of a considerable amount of sequence data, reconstructions are not always accurate and can result in incongruent topologies (Philippe et al., 2011). These limitations are partly due to methodological artifacts such as sequence misalignment (different software gives significantly different alignments (Wong et al., 2008)), false-orthologous gene assignment (due to horizontal transfer, gene duplication/loss events (Bapteste et al., 2004)) and homoplasy inherent to the data. These limitations prompted phylogeneticists to explore different types of signal representing rare genomic changes, such as intron indels, retroposon integrations, changes in organelle gene order, gene duplications and genetic code variants (Rokas and Holland, 2000). Although these genomic changes can be useful to validate some topological uncertainties, they have never been used to reconstruct complete phylogenies at the exception of the coherent mitochondrial phylogeny based on gene composition and gene order of mitochondrial genomes (Sankoff et al., 1992). This result offered, for the first time, a strong validation of the hypothesis that the macrostructure of mitochondrial genomes contains quantitatively meaningful information for phylogenetic reconstruction.

Gene order along nuclear chromosomes follows different evolutionary trends than along mitochondrial genomes (Burger et al., 2003), and it has been observed in several occasions that it comprises useful evolutionary information for phylogenetic reconstruction (Boore, 2006; Fertin, 2009). Many methods aiming at exploiting this trait as phylogenetic signal have been developed. They all belong to one of the four classical methodological categories *i.e.* the distance-based methods (Moret et al., 2001a; Wang et al., 2006; Guyon et al., 2009; Luo et al., 2012; Lin et al., 2012), the maximum-parsimony-based methods (Sankoff and Blanchette, 1998; Cosner et al., 2000; Moret et al., 2001b; Bourque and Pevzner, 2002; Tang and Moret, 2003; Bergeron et al., 2004; Zheng and Sankoff, 2011; Xu et al., 2011), the maximum-likelihood-based methods (Larget et al., 2005; Hu et al., 2011; Lin et al., 2013; Feng, 2017) and the quartet-based methods (Liu et al., 2005). Whether they are applied to sequences or to gene orders, these methodological categories harbor a variety of intrinsic limitations: computational complexity, sensitivity to short and long branch attraction (Felsenstein, 1978), requirement for good evolutionary models (Yang and Rannala, 2012), etc. Moreover, gene order-based methods were so far mainly applied to small bacterial or organelle genomes or to highly colinear genomes. The first phylogenetic reconstruction of eukaryotic nuclear genomes harboring different gene contents and different levels of synteny conservation was applied to the very large evolutionary span covered by the super-group of Unikonts and did not assess the performance of the method at known difficult branching positions such as the position of Rodentia relative to Primates and Laurasiatheria (Xu et al., 2011), or the position of *Candida glabrata* in Saccharomycetaceae (Lin et al., 2013; Hu et al., 2014). A recent improvement of this method taking into account balanced rearrangements, insertions, deletions, and duplications into an evolutionary model based on the principle of Double Cut and Join was applied to the phylogenetic reconstruction of 20 yeast species. It achieved accurate phylogeny reconstruction although the tree topology showed a couple of disagreements with previously published phylogenies (Feng, 2017).

We developed PhyChro with the aim of making the most of the evolutionary information derived from chromosome rearrangements. PhyChro is applied to 13 vertebrate and 21 Saccharomycotina yeast genomes and it reconstructs very accurate tree topologies even at known difficult branching positions.

## New approaches

PhyChro is a method for phylogenetic reconstruction based on synteny block and gene adjacencies. It relies on two important specificities. First, it uses synteny block adjacencies computed for all possible pairwise combinations of species instead of using synteny blocks universally shared by all the species involved in the reconstruction. This pairwise approach has the advantage to efficiently compare genomes with different levels of synteny conservation, without losing the wealth of synteny information that is shared by most closely related genomes. Second, PhyChro achieves tree reconstruction using the idea that for each synteny breakpoint, (a subset of the) genomes can be split into two disjoint groups depending on whether they support one block adjacency defining the breakpoint or the other. Formally, PhyChro relies on partial splits (Semple & Steel, 2001; Huson et al., 2004; Huber et al., 2005) (**Figure 1**), a generalization of the notion of split used in quartet-based methods. By exploiting partial splits associated to all identified breakpoints, PhyChro defines a distance between genome pairs, called *Partial Split Distance* (PSD), by counting the number of times that two genomes belong to different subsets of a partial split. Note that PSD is a measure defined on a set of *n* genomes contrary to other previously introduced distance measures based on the comparison of only two genomes at a time. Based on PSD, PhyChro reconstructs tree topologies with a bottom-up approach, by iteratively identifying those sister genomes that minimize the number of times they belong to different subsets of a partial split.

**Figure 1.**
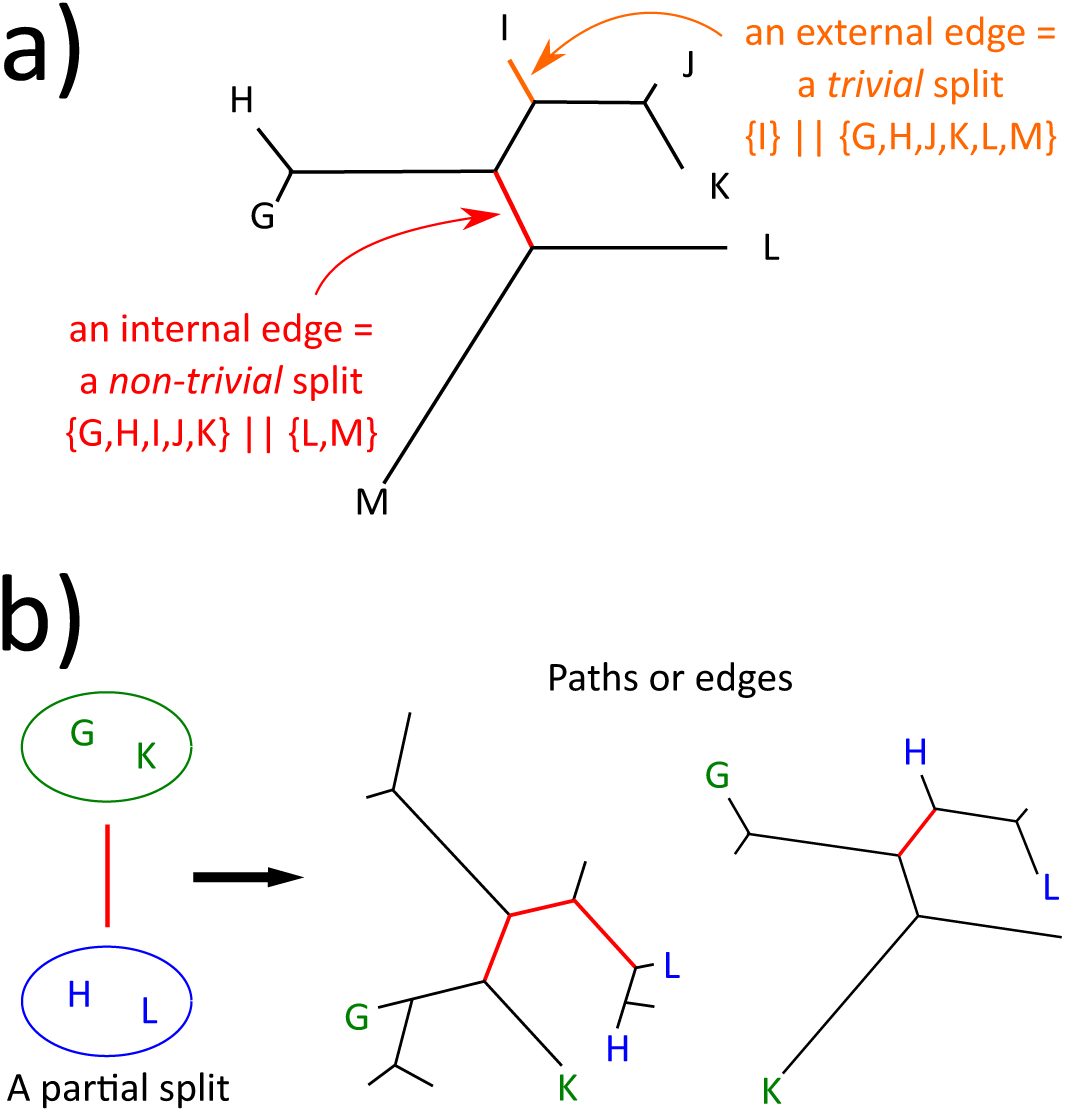
Splits and partial splits. **a)** Examples of trivial (orange edge) and non-trivial (red edge) splits. **b)** The two sets of genomes {*GK*} and {*HL*}, forming a partial split, uniquely determine a path (in red, in the left tree) or an edge (in red, in the right tree) that join the smallest subtrees including *G, K*, and *H, L*.

Intuitively, sister genomes are pairs of genomes sharing a high number of gene adjacencies at breakpoint positions. One can think to these pairs of genomes as being located close to each other but also as being located further away from all other genomes. Based on these intuitions, PhyChro (i) focuses on chromosomal rearrangement events supporting internal branches (useful for topology reconstruction) while ignoring all events that occurred on external branches (of no use for topology reconstruction), and (ii) minimises the differences between sister genomes, that is genomes separated by no internal branch, rather than maximising their similarity.

Contrary to distance-based methods, each pairwise distance depends on all genomes (as it depends on breakpoints identified through all genome comparisons) and, at each iteration, PhyChro recomputes distances from scratch between all pairs of genomes not yet included in the reconstruction. This iterated updating, affecting all entries of the distance matrix, is original to PhyChro and absent in distance-based methods. The Neighbor Joining (NJ) algorithm encodes the somewhat similar idea that pairs of genomes need not only be close to each other but also be distant from all others to be considered first in the reconstruction. This second condition is explicitly handled by the NJ algorithm, while PhyChro encodes it directly in its definition of genome distance. In conclusion, PhyChro is an algorithm whose basic data structure is the partial split and whose computational model is a bottom-up iterative reconstruction of the tree based on genome distances. These distances are computed by successive approximations, after the iterative elimination of inconsistencies in the set of partial splits.

PhyChro provides estimations for branch length and branch robustness. Extensive details on the algorithm and on the notions on which it relies are provided in the Materials and Methods section.

## Results

### Phylogenetic reconstruction of yeast and vertebrate species

We tested PhyChro on two different sets of species comprising 21 yeast and 13 vertebrate genomes. They harbor very different genome characteristics (in terms of genome size, number and density of genes, etc) as well as very different modes of chromosome evolution (number and rates of rearrangements, proportions of inversions versus translocations, whole genome duplication events, etc) (Drillon and Fischer, 2011). Previous analyses using the global level of divergence of orthologous proteins revealed that the evolutionary range covered by the Saccharomycotina subphylum exceeds that of vertebrates and is similar to the span covered by the entire phylum of Chordata (Dujon, 2006). Moreover, for both clades, the level of synteny conservation is highly variable between subclades with only 50% of genes belonging to synteny blocks between Amniota and fishes, or between yeast species from the Protoploid and CUG clades, while more than 95% of genes are conserved in synteny between Primates or between closely related species within the CUG clade (Drillon and Fischer, 2011). Finally, phylogenetic reconstructions in these two groups of species contain some ambiguous branching positions (sometimes controversial in the past), such as the position of Rodentia in the vertebrate tree or the position of *Candida glabrata* in the Saccharomycetaceae family of yeast, that we were interested to test with PhyChro.

We applied PhyChro on the sets of synteny blocks reconstructed with SynChro (Drillon et al., 2013, 2014) (see Methods) that resulted from genome pairwise comparison of the two sets of vertebrate and yeast species. The resulting tree topologies were compared to the reconstructions obtained with existing methods based on protein sequence comparisons, including PhyML, a maximum-likelihood-based method, ProtPars, a maximum-parsimony-based method, and Neighbor, a distance-based method (Guindon and Gascuel, 2003; Felsenstein, 1989) (see Methods).

The tree topology reconstructed by PhyChro for the 13 vertebrate species (**Figure 2a**) is identical to the topology produced by PhyML on 389 families of orthologs (illustrated in **Figure S1**). The position of Rodentia is correctly located, closer to Primates than to the Laurasiatheria. By comparison, ProtPars and Neighbor do not correctly place Rodentia (**Figure S1**). It should be noticed that PhyChro succeeded in correctly placing the rodent branch in the tree despite the fact that no partial split supports the existence of the branch splitting Primates and Rodentia from the other species. This is due to the fact that PhyChro, contrary to the other methods, does not construct the tree by identifying well supporting branches; rather, it avoids creating branches that are contradicted. This strategy allows PhyChro to treat difficult cases generated by small branches and characterised by very few rearrangements. In the specific reconstruction of Rodentia positioning, the detection of the short branch preceding their splitting with Primates, is rendered even more difficult by the important evolutionary history of Rodentia that likely erased the traces of the plausibly few ancestral rearrangements of Primates and Rodentia (see long branches in **Figure 2a**). PhyChro corresponding branch length equals zero and its confidence score *cS*, which assesses the robustness of the branch, is close to 0 (0.03, **Figure 2a**).

**Figure 2.**
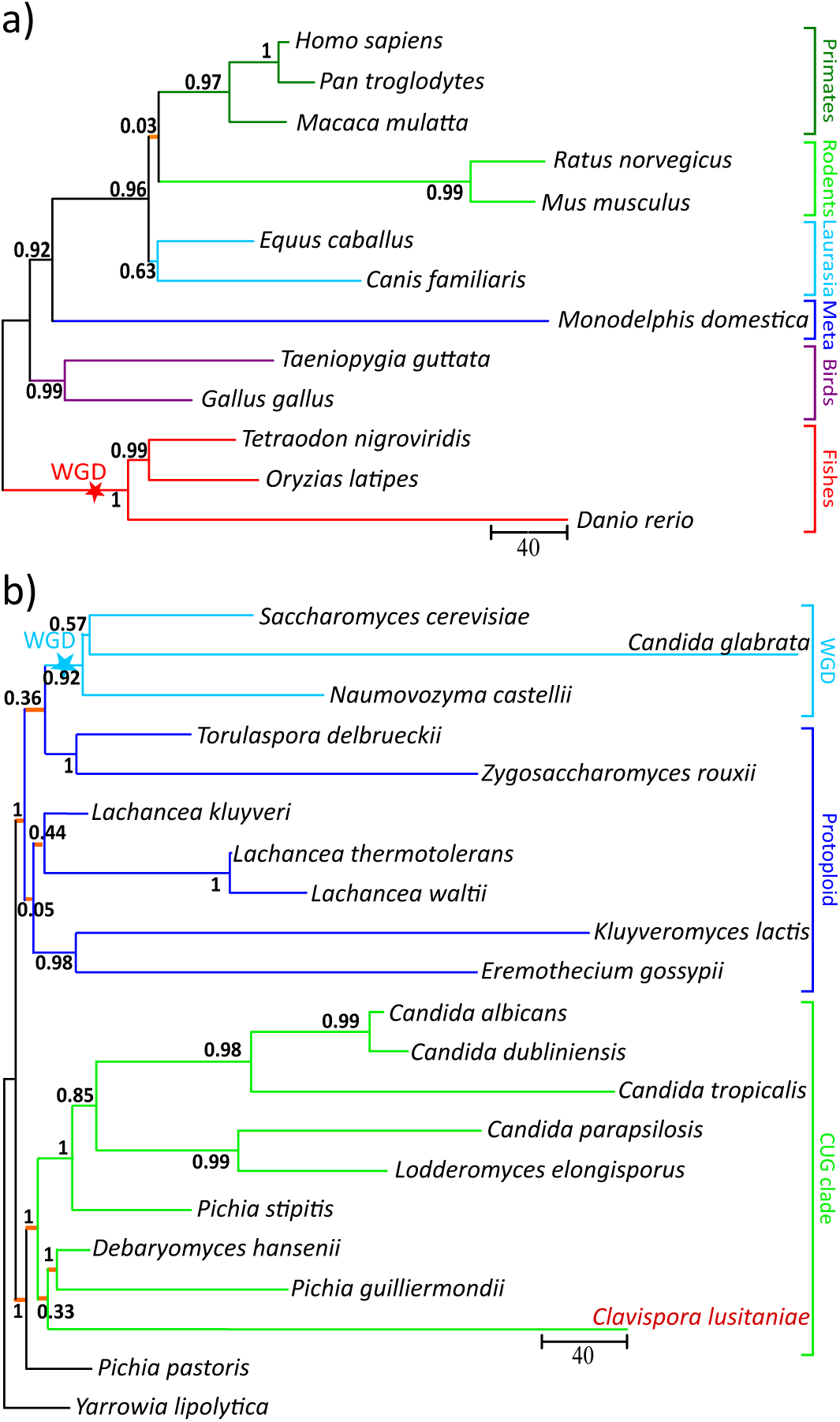
Phylogenies obtained with PhyChro for 13 vertebrate (a) and 21 yeast (b) species. Confidence scores that range between 0 and 1 are indicated on internal branches. Scale bars provide an estimation of the branch lengths, which correspond to the number of breakpoints, indirectly representing a number of rearrangements. For sake of clarity, internal branches with length smaller than 1 unit are represented in orange with an arbitrary small, but visible, length. Whole Genome Duplication events (WGD) are reported. *Clavispora lusitaniae* location in the tree is dubious and highlighted in dark orange.

In several ways, the tree topology reconstructed by PhyChro for the 21 yeast species is more accurate than the topologies obtained with either one of the 3 phylogenetic reconstruction methods, based on protein sequence comparison (**Figure 2b** and **Figure S1**). The first difference concerns the position of *Candida glabrata* relatively to *Saccharomyces cerevisiae* and *Naumovozyma castellii* (formely known as *Saccharomyces castellii*). It is known that phylogenies based on protein sequence analysis tend to artefactually place *C. glabrata* outside from *N. castellii* and *S. cerevisiae* (Kurtzman and Robnett, 2003; Hittinger et al., 2004) due to the short/long branch attraction problem (**Figure S1**). Previous studies based on shared patterns of gene losses and rearrangements showed that in fact, *N. castelli* is an outgroup to a clade containing *S. cerevisiae* and *C. glabrata* (Scannell et al., 2006; Gordon et al., 2009). Using the same macro-organisational information, PhyChro correctly recapitulates the phylogeny for these 3 species, despite the very long terminal branch length leading to *C. glabrata* present in its tree (**Figure 2b**). It should be considered that PhyChro reconstruction is automatic while the two previous ancestral gene ordering reconstructions have been manually derived.

In addition, note that PhyML erroneously locates *Pichia pastoris* as an outgroup while *P. pastoris* correctly branches at the root of the CUG clade according to PhyChro, Neighbor and ProtPars. Neighbor erroneously locates *Pichia stipitis* as a sister genome of *Debaryomyces hansenii* while *P. stipitis* is correctly positioned by PhyChro, PhyML and ProtPars. Concerning ProtPars, it erroneously splits the clade containing *Kluyveromyces lactis* and *Eremothecium gossypii* while the clade is correctly reconstructed by PhyChro, PhyML and Neighbor (**Figure 2b** and **Figure S1**). In all these instances, PhyChro outperforms the 3 classical methods based on protein sequence comparison.

The only topological uncertainty that remains corresponds to the position of *Clavispora lusitaniae*. According to PhyChro, this species branches as a sister genome to the clade containing *D. hansenii* and *Pichia guillermondii* (**Figure 2**), while according to PhyML and ProtPars, *C. lusitaniae* branches at the root of the CUG clade. Moreover Neighbor produces a third topology in this region of the tree (**Figure S1**). The confidence scores of the *C. lusitaniae* branch given by PhyChro, PhyML and ProtPars show uncertainties (0.33, 0.96 and 0.97, respectively) demonstrating that the topology associated to this branch remains doubtful.

Branch length estimates provided by PhyChro give interesting information notably for sub-clades where the synteny conservation is still high. For instance, the terminal branch length leading to the yeast *Lachancea thermotolerans* is computed to be very close to zero (0.33) showing that at most 1 rearrangement (larger than a six genes inversion) occurred in this genome since its divergence from its last common ancestor with *Lachancea waltii*, while long branches such as the ones leading to *C. glabrata, Danio rerio* or to Rodentia indicate the accumulation of a large number of chromosomal rearrangements. Note that branch lengths are under estimated for very distant genomes such as *Yarrowia lipolytica* and *P. pastoris* (as they are involved in very few partial splits).

### Comparison with MLGO, a gene-order based method for phylogenetic reconstruction

Currently, the only large-scale method to reconstruct gene order phylogenies is Maximum Likelihood for Gene Order Analysis (MLGO) (Lin et al., 2013). The two MLGO trees, issued from the same set of vertebrates and yeasts that we considered, are reported in **Figure S2**. These trees comprise a number of erroneous splits compared to the reference trees. We count two erroneous splits for vertebrates and seven for yeasts, contrary to PhyChro that reconstructs correctly both trees. For vertebrates the errors are due to the misplacements of *M. domestica* and Rodentia. For yeasts, *P. pastoris* is erroneously located closer to the Protoploid clade than to the CUG clade, *L. waltii* and the sister genomes *Torulaspora delbruechii* and *Zygosaccharomyces rouxii* are erroneously located in the Protoploid clade, and finally, *P. stipitis* is erroneously located in the CUG clade. As for PhyChro, *S. cerevisiae* and *C. glabrata* are correctly located.

### Robustness of PhyChro

#### Robustness of PhyChro on different definitions of synteny blocks

To test the sensitivity of PhyChro to different definitions of synteny block, we generated two sets of synteny blocks by using SynChro (Drillon et al., 2014) and i-ADHoRe 3.0 (Proost et al., 2012) and produced the corresponding trees for vertebrate and yeast species. On vertebrates, PhyChro based on i-ADHoRe synteny blocks gives a tree with an erroneous split corresponding to the misplacement of Rodentia (see **Figure S3a**). On yeasts, we count five erroneous splits in the tree reconstruction (**Figure S4a**). These discrepancies are explained by the lower proportion of genomes recovered in the synteny blocks generated by i-ADHoRe than by SynChro, as illustrated in **Figures S3bc** and **S4bc**. A global comparison of block size distributions generated by i-ADHoRe and SynChro over all pairwise comparisons between vertebrate and yeast genomes, is reported in **Figure S5**. We observe that SynChro allows for small blocks made of only 2 genes (noted also in (Drillon et al., 2014)) while i-ADHoRe only allows blocks of at least 3 genes, and that the number of small blocks (< 21 genes) produced by SynChro is systematically larger than for i-ADHoRE. For pairs of genomes that underwent many rearrangements and, in consequence, would have a low synteny conservation, the small blocks detected by SynChro are expected to play a crucial role. This is visually observable in the matrices of **Figures S3bc** and **S4bc** showing higher synteny coverage (lighter blue and darker red colours) for SynChro than for i-ADHoRE for all species pairs. On the other hand, one observes that i-ADHoRE generates a greater number of large blocks (≥ 21) than SynChro (**Figure S5**). This ensures that for pairs of genomes for which synteny blocks allow for more than 60% coverage, SynChro and i-ADHoRE show comparable success, as illustrated by the red coloured cells in the matrices of **Figures S3bc** and **S4bc**. In conclusion, a better synteny coverage reached for all pairs of species allows PhyChro to perform better on SynChro than on i-ADHoRE blocks.

It is also interesting to note that modulating the size of micro-rearrangements tolerated within synteny blocks with the Δ parameter from SynChro (bigger the Δ, larger the micro-rearrangements tolerated) has an effect on the number of partial splits contradicting a given topology. For example, PhyChro run with Δ = 3 (by default, see Methods) finds 36, 37 and 42 partial splits that contradict the ((Primates, Rodentia), Laurasiatheria), (Primates, (Rodentia, Laurasiatheria)) and ((Primates, Laurasiatheria), Rodent) topologies, respectively. By augmenting Δ to 4 (that is, being more tolerant for larger micro-rearrangements within synteny blocks), PhyChro finds 24, 37 and 53 contradictory partial splits, respectively. These numbers provide confidence in the ((Primates, Rodentia), Laurasiatheria) topology and, since none of the topologies has zero contradictions, they also show that homoplasy is present.

#### Robustness of the algorithm with respect to simulated genomes

In order to test PhyChro on a large set of simulated data representative of yeast and vertebrate genomes, we used computer simulations based on a realistic evolutionary model. We started with hypothetical ancestral genomes characterized by 5,000 genes distributed along 8 chromosomes for yeasts and by 18,000 genes distributed on 23 chromosomes for vertebrates. In both cases, we simulated random tree topologies with 21 leaves for yeasts and 13 leaves for vertebrates. The method for the construction of a random tree takes genes as building blocks and goes as follows:

1. it generates a random binary tree by defining the branching nodes uniformly over the time scale, with the exception of the first branching which is put at the root. More precisely, for each branching, it selects a leaf to split. It does it by going recursively from the root to the leaf by passing through internal nodes of the tree, with a half probability of choosing the right or the left subtree at an internal node. Once it selects a leaf to split, it attaches to it two new leaves. This construction is repeated until the number of leaves is equal to the expected number of species (21 species for yeasts and 13 for vertebrates).
2. based on the tree produced in step 1, it simulates chromosomal rearrangements along each branch of the tree, following a Poisson distribution, such that the average number of events from the ancestor (located at the root) to the species (located at the leaves) is approximately 500 for yeasts and 1000 for vertebrates. (We note that these values are comparable to those obtained on actual yeast and vertebrate genomes (Drillon and Fischer, 2011)). Rearrangements were distributed on the tree according to the following proportions: 60% of inversions, 29.79% of reciprocal translocations, 5% of duplications, 5% of deletions, 0.1% of fusions, 0.1% of fissions, 0.01% of whole genome duplications (WGD) (Ma et al., 2006; Drillon and Fischer, 2011). Following a WGD event, one of the two copies of each duplicated gene was deleted with a probability of 80% (Wolfe and Shields, 1997). The number of genes involved in an inversion, duplication and deletion was chosen following a Poisson distribution (where the parameter of the distribution was set to 5 genes for inversions and duplications, and to 1 gene for deletions).

The simulated genomes produced by this approach are consistent with actual yeast and vertebrate genomes in terms of number of genes, number of chromosomes and number of rearrangements along the branches of the trees. For the analysis, the minimum number of rearrangements per branch was set to 1 or to 10 for both yeast and vertebrate trees, and 100 simulations were generated in each case. Synteny blocks were computed between all pairs of simulated genomes (note that here genes are represented by numerical identifiers, not by actual nucleotide or amino-acid sequences) and PhyChro was run on these simulated genomes to compare the predicted topologies with the known (simulated) ones. For determining PhyChro success rate, we counted the number of splits in the trees that were correctly and incorrectly reconstructed by PhyChro. For a minimum number of rearrangements per branch set to 1, the results are reported in **Figure 3**, where one observes that PhyChro is able to reconstruct correct tree topologies without any erroneous split in 79% of the cases for vertebrates and 61% for yeasts, and for the incorrect ones, in most cases (17% for vertebrates and 30% for yeasts), we record just one incorrect split per tree. Over all trees, 97% of the splits are correctly predicted by PhyChro, both for vertebrates and yeasts, and, most importantly, incorrect splits mainly correspond to very short branches, that is branches where only very few rearrangement events took place (see inset plot in **Figure 3**). If we set the number of events in a branch to be at least 10, the number of correct trees for vertebrates increases to 86% and for yeasts to 69%, with 98% and 97% of the splits that are correct over all trees, for vertebrates and yeasts, respectively.

**Figure 3.**
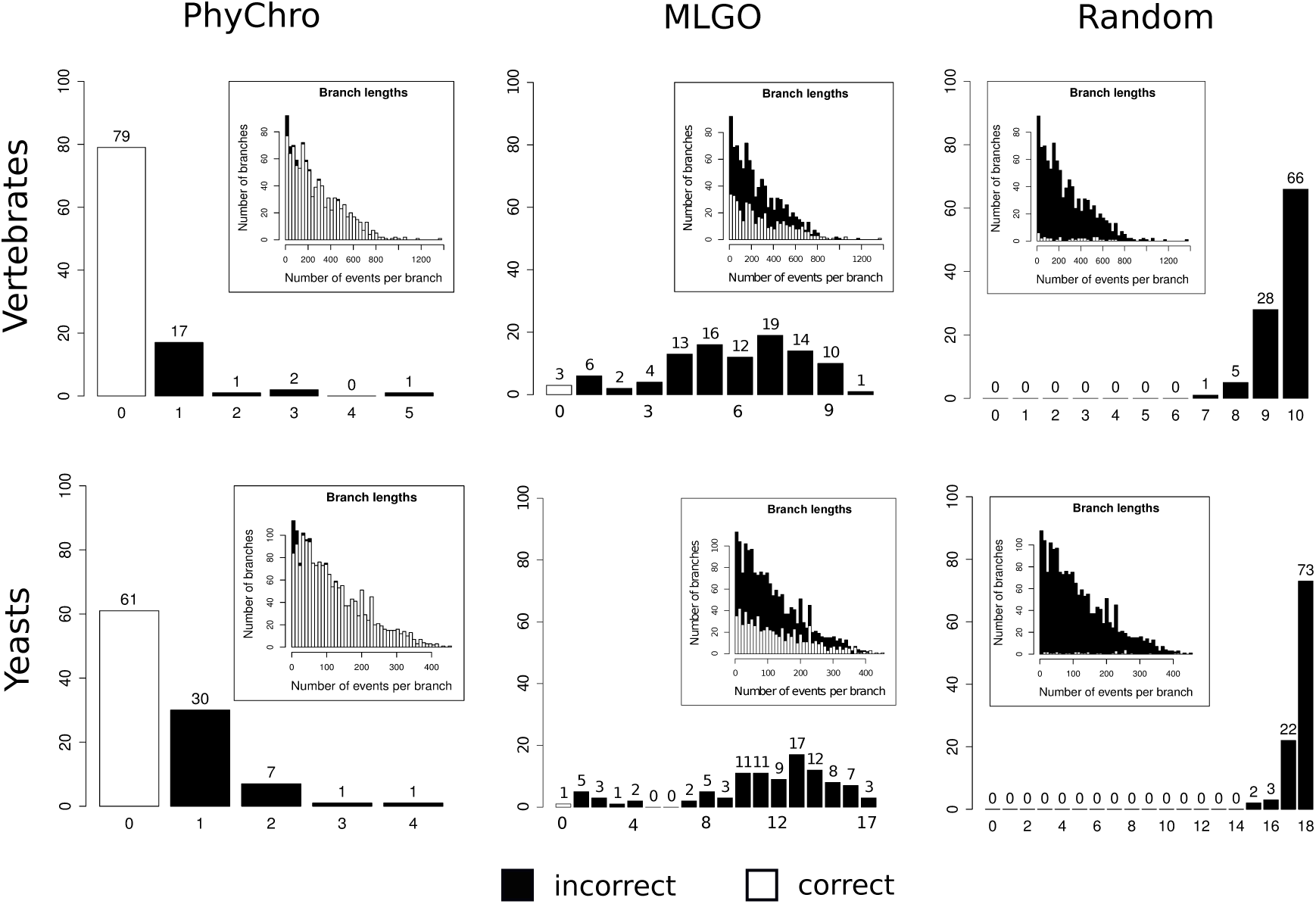
PhyChro, MLGO and random reconstructions tested on simulated trees. Simulated phylogenetic trees describing rearrangement events were generated for vertebrate-like (top) and yeast-like (bottom) genomes and used to check whether PhyChro (left) and MLGO (center) could correctly reconstruct the original phylogeny from the corresponding sets of simulated genomes. The simulated trees have been used also to check to which extent a random assignment of rearrangements (right) on the branches could correctly reconstruct the original phylogeny from the corresponding sets of simulated genomes. The histograms report the number of trees with a fixed number of incorrect splits predicted by the three methods. The inset plots represent the distribution of the number of branches with a fixed length (corresponding to a number of simulated rearrangements that were applied to these branches) in the simulated trees, and describe how many of those branches have been reconstructed correctly (white) or incorrectly (black) by a method.

This analysis helps to evaluate a confidence threshold for scores *cS*. In fact, 99% of correct splits are obtained with a score *cS* ≥ 0.2 for the 100 simulated genomes for yeasts, and with a score *cS* ≥ 0.6 for the 100 simulated genomes for vertebrates. This means that in the yeast phylogenetic tree reconstructed by PhyChro, the only weakly supported branch (scoring 0.05) is the one locating *E. gossipii* and *K. lactis* within the Protoploid clade, while the branch locating *C. lusitaniae*, displays a sufficiently strong *cS* score (0.33) to be trusted (**Figure 1b**). For vertebrates, as discussed above, the position of Rodentia in the tree remains very weakly supported (**Figure 1a**).

A random shuffling of species in the 100 randomly generated trees is reported in **Figure S3**, where we note a shape of the distribution of errors that has a complementary tendency compared to the one obtained for PhyChro, that is the vast majority of events associated to a branch is incorrect and the number of erroneous splits corresponds, most of the times, to the number of internal branches (10 for vertebrates and 18 for yeasts). This corresponds to no correct trees obtained for both vertebrates and yeasts; we note that only the 1% of the splits are correct for yeasts and only the 3% for vertebrates. The same test, based on the same dataset of trees (and the same synteny blocks considered by PhyChro and the random tree analyses), has been realized on MLGO (**Figure S3**). MLGO works much better than the random case but yet is far from PhyChro performance: 3% of trees are correct for vertebrates and 1% for yeasts. Many of the trees that are reconstructed by MLGO have a high number of erroneous splits (57% for yeast and 42% for vertebrates) both for vertebrates and yeasts.

#### Adding new genomes to PhyChro reconstructions - a case study

The arrival of new sequenced genomes asks for their integration in the phylogenetic tree, and PhyChro can be profitably used to insert these new species. As an example, we considered the vertebrate tree in which, some of the species are known to be difficult to handle. In this respect, the literature contains an open debate because mammalian species positioning appears sensitive to the evolutionary information taken into consideration in phylogenetic reconstruction (Romiguier et al., 2013). We added three recently sequenced genomes, the cow, the pig and the lizard. PhyChro tree reconstruction (**Figure 2a**) correctly placed *Anolis*, the lizard, close to the birds; both are known to be members of Diapsida. It also added *Bos taurus* (cow) and *Sus scrofa* (pig) to the clade including horse and dog, with the nesting (horse, (dog, (cow, pig))). See **Figure S6** for an illustration of the resulting tree, and its legend for an analysis of the dubious position of the cow and the pig with respect to the horse and the dog (this reconstruction being of interest in exemplifying limits and power of PhyChro).

## Discussion

### PhyChro, a new strategy of phylogenetic recontruction

An important effort was made in this work to identify how chromosomal breaks coming from chromosomal rearrangements could be used as phylogenetically informative characters to perform phylogenetic reconstructions. PhyChro differs in a fundamental way from the classical reconstruction methods. The first difference comes from the pairwise comparison approach between genomes which allows us to make the most out of the synteny information shared between closely and distantly related genomes at the same time. Another difference comes from the definition of 2 functions (f_inc_ and f_comp_, see Methods) which represent, respectively, the number of times where two genomes are split in two groups of incompatible adjacencies and the number of times where they are grouped together (not split) based on shared adjacencies. The ratio between these two functions is used to identify the least incompatible pairs of species from which sister genomes will be defined. The main originality of PhyChro is that it identifies sister genomes by minimizing the number of incompatible adjacencies rather than by maximizing the number of shared rearrangements. Formally, PhyChro bases its tree reconstruction on the Partial Split Distance. This distance relies on the notion of partial split that allows to record the number of incompatible adjacencies for pairs of genomes among a set of genomes. Hence, PhyChro does not try to combine internal branches into a tree topology, but rather it reconstructs the topology by iteratively identifying genomes and ancestral genomes that are closely related. It uses a bottom-up approach, similarly to what is done in distance-based methods. Note that PSD is a measure defined on a set of *n* genomes contrary to other previously introduced notions, measuring genome rearrangements, that are based on the comparison of only pairs of genomes. An example is the well known Breakpoint Distance (BD), defined to be the number of breakpoints observable from the comparison between two genomes. The notion was first used in (Nadeau and Taylor, 1984), then formally defined for one (Watterson et al., 1982; Sankoff and Blanchette, 1997) and multiple (Pevzner and Tesler, 2003; Tannier et al., 2009) chromosomes. The direct comparison between PSD and BD is impossible given that for two genomes *G, H* among *n*, the distance *BD*(*G, H*) depends only on *G, H* while *PSD*(*G, H*) depends on the *n* genomes. When reconstructing phylogenies, knowledge on the way pairs of genomes split in the tree (recall that the notion of non-trivial split is based on at least four genomes and not on pairs nor triplets) is primordial and one can only gather it through comparisons between all genomes involved in the reconstruction. This is why the intrinsic nature of a measure based on *n* genomes, like PSD, is expected to bring fundamental information for phylogenetic tree reconstruction. It is important to notice that PSD counts only those breakpoints that are supported by at least a quadruplet of genomes, and associated to rearrangements shared by at least two genomes, while BD counts all breakpoint events including those associated to rearrangements that are specific to a given genome (occurring on the external branches of a tree).

Thanks to this reconstruction strategy, PhyChro is less affected by “short-branch” attraction, which often leads distance-based methods to put genomes having undergone a lot of rearrangements/mutations higher in the tree than they belong. Another originality of PhyChro is that it provides branch length estimates that reflect the level of chromosome plasticity rather than the rates of punctual mutations, as all classical methods of phylogeny reconstruction do. In addition, PhyChro allows estimation of the robustness of branches in a way that is radically different from the bootstrap methods. The advantage here is that computing confidence scores is very fast as it does not involve additional tree reconstructions.

### Phylogenetic reconstruction based on chromosomal rearrangements

We showed through the analysis of simulated genomes that PhyChro generates very accurate tree topologies by successfully reconstructing known tree topologies. Applications of PhyChro to real biological datasets comprising different types of genomes (yeasts and vertebrates) and covering different evolutionary ranges shows that chromosomal rearrangements are indeed phylogenetically informative and that accurate phylogenies can be reconstructed solely based on these large scale mutational events. This success demonstrates that the evolutionary signal that derives from chromosome rearrangements comprises at least as much phylogenetic information as the local information present in protein sequences. Moreover, we showed that PhyChro reconstructions are at least as accurate as the best reconstructions deriving from classical methods that use protein sequence comparisons. We also show that at particularly difficult branching positions, such as that of *C. glabrata* relatively to *S. cerevisiae* and *N. castellii*, PhyChro outcompetes all other methods of phylogenetic reconstruction.

Another important application of PhyChro was realized (with the same parameters used for vertebrates and yeasts species) on scleractinian corals, the foundation species of the coral-reef ecosystem. Corallimorpharians had been proposed to originate from a complex scleractinian ancestor that lost the ability to calcify in response to increasing ocean acidification, suggesting the possibility for corals to lose and gain the ability to calcify in response to increasing ocean acidification. A phylogenetic analysis based on 1 421 single-copy orthologs combined with PhyChro phylogenetic reconstruction allowed to disprove this hypothesis contributing evidence for the monophyly of scleractinian corals and the rejection of corallimorpharians as descendants of a complex coral ancestor (Wang et al., 2017).

These results suggest that synteny information should be integrated more broadly in future phylogenetic reconstruction analysis pipelines.

## Materials and Methods

The classical notions of synteny blocks, breakpoints, splits and partial splits are recalled. We introduce the notions of “Partial splits associated to breakpoints” and of “Partial Split Distance” that are central in PhyChro.

### Synteny blocks

A pairwise genome comparison *G/H* (or equivalently *H/G*), between the two genomes *G, H*, identifies chromosomal segments with conserved orthologous gene order. These segments are called *synteny blocks*, and are also referred to as *blocks*. Without loss of generality, we call *B* both the occurrences of the synteny block *B* in *G* and in *H*. Different definitions of synteny blocks have been proposed before (Ferretti et al., 1996; Roedelsperger and Dieterich, 2010; Pham and Pevzner, 2010; Proost et al., 2012; Drillon et al., 2014) and they are based on different conditions on the proximity between orthologs. PhyChro works with blocks *B* that verify the following five conditions:

- *B* is described by its pairs of homologous genes in *G* and *H*, called *anchors* for *B*. Since a gene can have several homologs, it can be involved in the definition of several anchors (within the same block or in different ones).
- the first and the last genes of *B* in *G* (*H*) have homologs in the corresponding block *B* in *H* (*G*). We say that *B* in *G* (*H*) is delimited by its first and last anchors.
- *B* is unique, in the sense that duplicated blocks are not explicitly handled and they are defined as independent blocks. For instance, if *B* is duplicated in *G* but not in *H*, the two copies of the block are considered as distinct in *G* and as overlapping in *H*.
- *B* is oriented or signed, and in particular, *B* can have a different orientation in *G* and in *H*. The orientation of *B* in a genome *G* maybe fixed in some arbitrary way or might depend on conditions that are specific to the definition of a block, such as the order and the orientation of its genes. When impossible to be established, a block orientation is left undetermined and the block is called “unoriented” or “unsigned”. The orientation of a block allows us to differentiate its right and left ends (in order to determine which of its extremities is involved in a breakpoint): the “end” of *B* corresponds to the “beginning” of −*B* and reciprocally.
- *B*, in *G* or *H*, can overlap or be included in another block.

A block *B* is called *telomeric* if it is the first or the last block of a chromosome in *G* or in *H*.

### Breakpoints

Chromosomal rearrangements generate synteny breakpoints, or analogously, synteny block adjacencies. Given a block *B* obtained through the comparison *G/H*, a breakpoint is defined by the pair [(*B A*)_*G*_, (*B C*)_*H*_] of block adjacencies (*B A*) in *G* and (*B C*) in *H*. In I of **Figure 4**, for instance, the right end of block *B* is contiguous to the left ends of blocks *A* and *C* in genomes *G* and *H*, respectively. Since blocks are oriented, notice that the same breakpoint might correspond to [(*B A*)_*G*_, (−*C* −*B*)_*H*_], where −*B* has −*C* on its left end instead. Notice also that synteny blocks derived from duplications or chromosome fusions/fissions do not generate pairs of block adjacencies and therefore are not explicitly considered here. Blocks derived from translocations, inversions and transpositions of DNA segments are the only ones that are informative in our analysis. Each block (except the telomeric ones) should, in theory, lead to two breakpoints (one at each end of the block, see I in **Figure 4**). However, complex gene-order configurations might lead to a reconstruction of synteny blocks that overlap, are included in one another or are unoriented (like for blocks reconstructed by SynChro (Drillon et al., 2014)). In the following, we consider as breakpoints only those pairs of regions in *G* and in *H* for which preceding and following blocks are unambiguously identified (and ignore the others).

**Figure 4.**
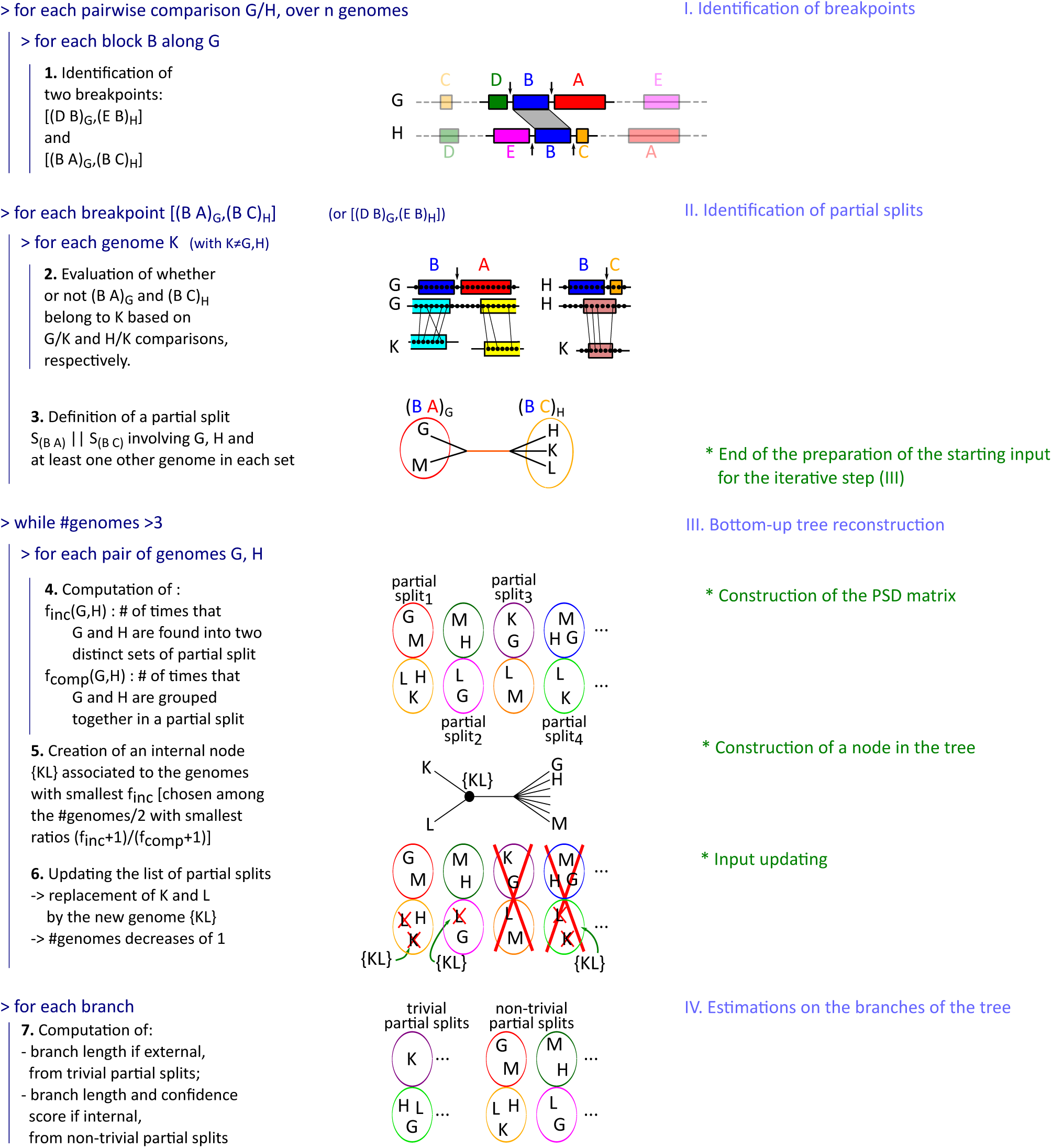
PhyChro algorithm. The four main parts and the seven steps, briefly described here, are detailed in the main text.

### Splits

A split is a bipartition of a set of taxa. **Figure 1a** illustrates an example of a split and of a trivial split, that is, a split induced by an external edge connecting a leaf to the rest of the tree. Splits play an important role in phylogenetic reconstruction (Bandelt and Dress, 1992; Huson et al., 2010) as each edge of an unrooted tree is univocally associated to a split. In fact, an edge splits taxa into the two disjoint subsets *S*_1_, *S*_2_ labeling the leaves of the subtrees rooted at the extremes of the edge. We note that the union of *S*_1_, *S*_2_ covers the full set of taxa. In evolutionary terms, we think of genomes in *S*_1_ (or *S*_2_) as having undergone a number of common ancestral rearrangements, specifically the ones that occurred along the edge, that genomes in *S*_2_ (*S*_1_) did not undergo. Strictly speaking, it cannot be established whether these rearrangements took place for *S*_1_ or for *S*_2_ because the tree is not rooted. Hence, ideally, for the reconstruction of a phylogenetic tree, one could hope (i) to recover rearrangements from genomic data, (ii) to define splits of genomes sharing the rearrangements and (iii) to reconstruct the edges of the tree by combining splits identified from the rearrangements.

### Partial splits

For the purpose of tree reconstruction, traces of chromosomal rearrangements may have disappeared in some genomes (due to the accumulation of other rearrangements), and it might become impossible to recover splits. This is why, we shall use a generalisation of the concept of split to the one of partial split. This notion was introduced in (Semple & Steel, 2001; Huson et al., 2004; Huber et al., 2005). Formally, a partial split is a pair of non-empty disjoint sets of taxa. Intuitively, given an unrooted phylogenetic tree whose leaves are labelled by different taxa and given some path *c* in the tree, we say that *c* induces a partial split of the sets of genomes *S*_1_, *S*_2_ if: 1. *S*_1_, *S*_2_ are constituted by some (possibly all) of the taxa associated to the subtrees rooted at the extremes of *c*; 2. in each *S*_*i*_, for *i* = 1, 2, there are at least two taxa that are connected by a shortest path passing through the root of the corresponding subtree (**Figure 1b**). We note that, by definition, *S*_1_ ∩ *S*_2_ = Ø and, also, that *S*_1_ ∪ *S*_2_ does not necessarily correspond to the full set of taxa in the subtrees rooted at the extremes of *c. A fortiori, S*_1_ ∪ *S*_2_ does not necessarily correspond to the full set of taxa in the complete tree, as it is the case for splits. In fact, a split is a partial split where *c* is an edge, but a partial split induced by an edge need not be a split because of condition 1. As for splits, we think of genomes in *S*_1_ (or *S*_2_) as having undergone a number of common ancestral rearrangements, specifically the ones that occurred along the path *c*, that genomes in *S*_2_ (*S*_1_) did not undergo.

As for splits, we say that a partial split is *trivial* when one of the two subsets *S*_1_, *S*_2_ is a singleton. Notice that trivial partial splits do not bring information on the topology of the tree (since the set of trivial partial splits is the same for all topologies) and are not used in tree reconstruction. We shall use them to estimate the length of the terminal branches though, that is, branches leading to leaves in the tree.

### Testing the conservation of block adjacencies

Given a breakpoint [(*B A*)_*G*_, (*B C*)_*H*_] in the comparison *G/H*, we test for the presence of (*B A*)_*G*_ in a genome *K* (by definition, (*B A*)_*G*_ ∈ *G*). The test is similarly stated for (*B C*)_*H*_. The test does not directly search for blocks *B* and *A* in *K* because they might not have direct equivalents in *G/K*. Instead, it infers the presence of the adjacency (*B A*)_*G*_ in *K* at the gene level, by testing whether the genes flanking the (*B A*)_*G*_ adjacency in *G*, that is, the right end of block *B* and the left end of block *A*, have syntenic homologs in *K*. More precisely, the test compares *G* and *K* and determines whether there is a synteny block *D* in *K* and *G* such that the following conditions are satisfied (we refer to the notation employed in **Figure 5** - see also **Figure S7**):

**Figure 5.**
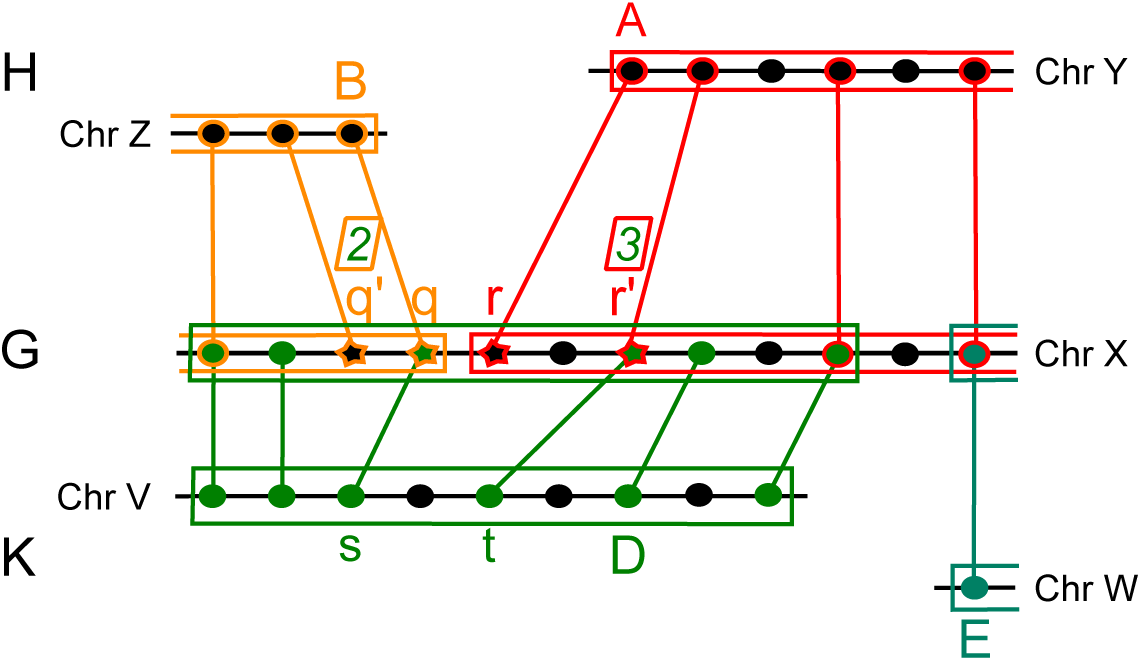
Localization of the adjacency (*B A*)_*G*_ in the genome *K*. **Conservation of the adjacency (*B A*)_*G*_ in the genome *K*.** Genes are indicated as dots or stars. Stars, in *G*, are used for the two last (*q*′ and *q*) and first (*r* and *r*′) anchors of blocks *B* and *A* in the comparison *G/H*. Red, yellow and green colors are used to highlight anchors associated to the blocks *A, B* and *D*, obtained in the comparisons *G/H, G/H* and *G/K*, respectively. Genes *q*′, *q, r* and *r*′ belong to the same block *D* in *G/K*. The number of anchors of *D* lying before *q*′ (after *r*′), and possibly including it, is indicated above *q*′ (*r*′) within a square. Gene *s* (*t*) is the anchor of *D* whose homolog in *G* lies in the right (left) most position of *B* (*A*). Homology is indicated by links among genes occurring in different genomes: *s* is homolog of *q* and *t* of *r*′. Note that, here, the three conditions (i)-(iii) discussed in the text are satisfied and that *K* ∈ (*B A*)_*G*_.

i. the two last anchors (or syntenic homologs) *q*′, *q* of *B* and the two first anchors *r, r*′ of *A*, along *G*, belong to the same synteny block *D* in *G/K*;
ii. *q*′, *r*′ are preceded and followed along *G*, respectively, by at least two other anchors in *D* (possibly including themselves).
iii. let *s* be the anchor of *D* in *K* whose homolog in *G* lies in the right most position of *B*, and let *t* be the anchor of *D* in *K* whose homolog in *G* lies in the left most position of *A*. Then, the sum of the number of genes between *s* and *t* in *K* and between their homologs in *G* (see **Figure 5**) is at most 4.

Conditions (i) and (ii) guarantee block *D* in *G* to overlap several anchors of *A* and *B* in *G*, and condition (iii) ensures the genes forming the (*B A*)_*G*_ adjacency in *G* and *K* to be in physical proximity. Such proximity is computed for a maximum of 4 genes between the two anchors *s* and *t* in **Figure 5**. All values from 3 to 6 have been tested to choose the best parameter for yeasts and vertebrates. (Note that value 3 is too strict and value 6 brings noise in the construction.) These three conditions introduce some flexibility in the definition of synteny conservation, without being too permissive. If they are all satisfied, we say that the adjacency belongs to *K* and write (*B A*)_*G*_ ∈ *K*. If *q* and *r* belong to the same block *D* in *G/K* but some of the conditions fail, we still say that (*B A*)_*G*_ ∈ *K* and consider the relation as *weakly* supported. These weak adjacencies can be due to false ortholog assignments or to small inversions. In all other cases, we say that (*B A*)_*G*_ ∉ *K*.

### Partial splits assigned to breakpoints

Given a breakpoint [(*B A*)_*G*_, (*B C*)_*H*_], we define a partial split by identifying two sets of genomes, *S*_(*BA*)_ and *S*_(*BC*)_, where *S*_(*BA*)_ comprises genomes sharing the adjacency (*B A*)_*G*_ and *S*_(*BC*)_ comprises genomes sharing the adjacency (*B C*)_*H*_. For this, we apply the above adjacency test, checking whether the adjacencies (*B A*)_*G*_ and (*B C*)_*H*_ derived from the *G/H* comparison are present in a genome *K* or not, for all *K* ≠ *G, H*. Namely, *K* ∈ *S*_(*BA*)_ if and only if (*B A*)_*G*_ ∈ *K*, and *K* ∈ *S*_(*BC*)_ if and only if (*B C*)_*H*_ ∈ *K*.

Notice that a genome *K* that neither contain (*B A*)_*G*_ nor (*B C*)_*H*_ belongs to none of the two sets. Also, a genome *K* may contain, at the same time, the two adjacencies defining a given breakpoint. This ambiguous case might occur either for a breakpoint [(*B A*)_*G*_, (*B C*)_*H*_] when *C* follows *A* in *G* and *A* is small enough to make condition (iii) true for (*BC*)_*H*_ in *K* (see **Figure S8a**), or for a breakpoint [(*B A*)_*G*_, (*B* − *A*)_*H*_] when *A* is small enough to make (*BA*)_*G*_ ∈ *K* and (*B* − *A*)_*H*_ ∈ *K* (see **Figure S8b**).

Intuitively, the coexistence of (*B A*)_*G*_ ∈ *K* and (*B C*)_*H*_ ∈ *K*, for some *K*, indicates that (*B A*)_*G*_ and (*B C*)_*H*_ are too “similar” to claim that they support a split. Therefore, it is only when the two sets of genomes *S*_(*B A*)_, *S*_(*B C*)_ are disjoint that we say that they form a *partial split*, denoted *S*_(*B A*)_1*S*_(*B C*)_, associated to the breakpoint [(*B A*)_*G*_, (*B C*)_*H*_] (**Figure 1b**).

### The Partial Split Distance

Given a set of genomes, genome pairwise distances can be computed by considering the set of partial splits associated to all breakpoints issued from all pairwise genome comparisons. For this, we shall define two functions, f_inc_ and f_comp_, on the list of non-trivial partial splits.

The first one, f_inc_(*G, H*) (where “inc” stands for incompatible), counts the number of times that genomes *G* and *H* belong to different subsets of a partial split (as for partial splits 1 and 2 in III of **Figure 4**):

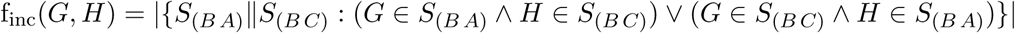

The second function, f_comp_(*G, H*) (where “comp” stands for compatible), counts the number of times that genomes *G* and *H* are found in the same subset of a partial split, *i.e.* sharing a same adjacency (as for the partial split 4 in III of **Figure 4**):

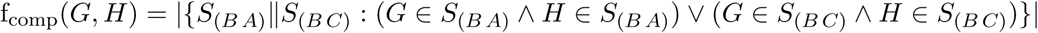

The function f_inc_ represents an “internal” distance between genomes, and we call it *Partial Split Distance*, PSD in short. Intuitively, given two genomes, PSD is proportional to the number of rearrangements that occur along the internal branches separating these two genomes in the phylogenetic tree that we want to reconstruct. The number of these rearrangements is estimated with *f*_*inc*_, by using the number of non-trivial splits separating the two genomes. This means that sister genomes, that is genomes separated by no internal branch, should have a PSD distance equal to zero (independently of the length of their external branches). This property will be used to identify sister genomes and to reconstruct phylogenies bottom-up (see below). In the same way, very close genomes, separated by few and short internal branches, should have a PSD close to zero. However, because *f*_*inc*_ is defined from non-trivial splits, very distant genomes, which do not share many adjacencies with other genomes and, therefore, are not involved in many non-trivial splits, have also a PSD very close to zero with all other genomes. To take into account this fact, we consider *f*_*comp*_ and use the ratio ℛ = (f_inc_ + 1)*/*(f_comp_ + 1) to discriminate among pairs of genomes that have a very small internal distance (f_inc_ close to zero) those that are very closely related (high f_comp_ value) from those that are very distantly related (f_comp_ close to zero).

Note that, if f_inc_(*G, H*) ≠ 0 then there exists at least one non-trivial partial split *S*_(*B A*)_‖*S*_(*B C*)_ that separates *G* from *H*. This means that there exist genomes *K, L* such that (*G, K* ∈ *S*_(*B A*)_ ∧ *H, L* ∈ *S*_(*B C*)_) ∨ (*G, K* ∈ *S*_(*B C*)_ ∧ *H, L* ∈ *S*_(*B A*)_). Ideally, this suggests that in a phylogenetic reconstruction involving genomes *G, H, K, L*, the two genomes *G, H* should not be considered as sister genomes. In reality, as mentioned above, it might be difficult to unravel complete information from breakpoints (due either to convergence or to the accumulation of rearrangements) and one might have to treat as sister genomes, those pairs of genomes that display the smallest f_inc_ value, even if it is different from 0.

### The PhyChro algorithm

Phylogenetic reconstruction based on partial splits is a more delicate problem than tree reconstruction based on splits (Bandelt and Dress, 1992; Huson et al., 2010; Semple & Steel, 2001; Huson et al., 2004; Huber et al., 2005). PhyChro comprises four main parts (I, II, III and IV; see **Figure 4** and **Table 1**) divided into 7 major steps that are detailed below.

**Table 1:**
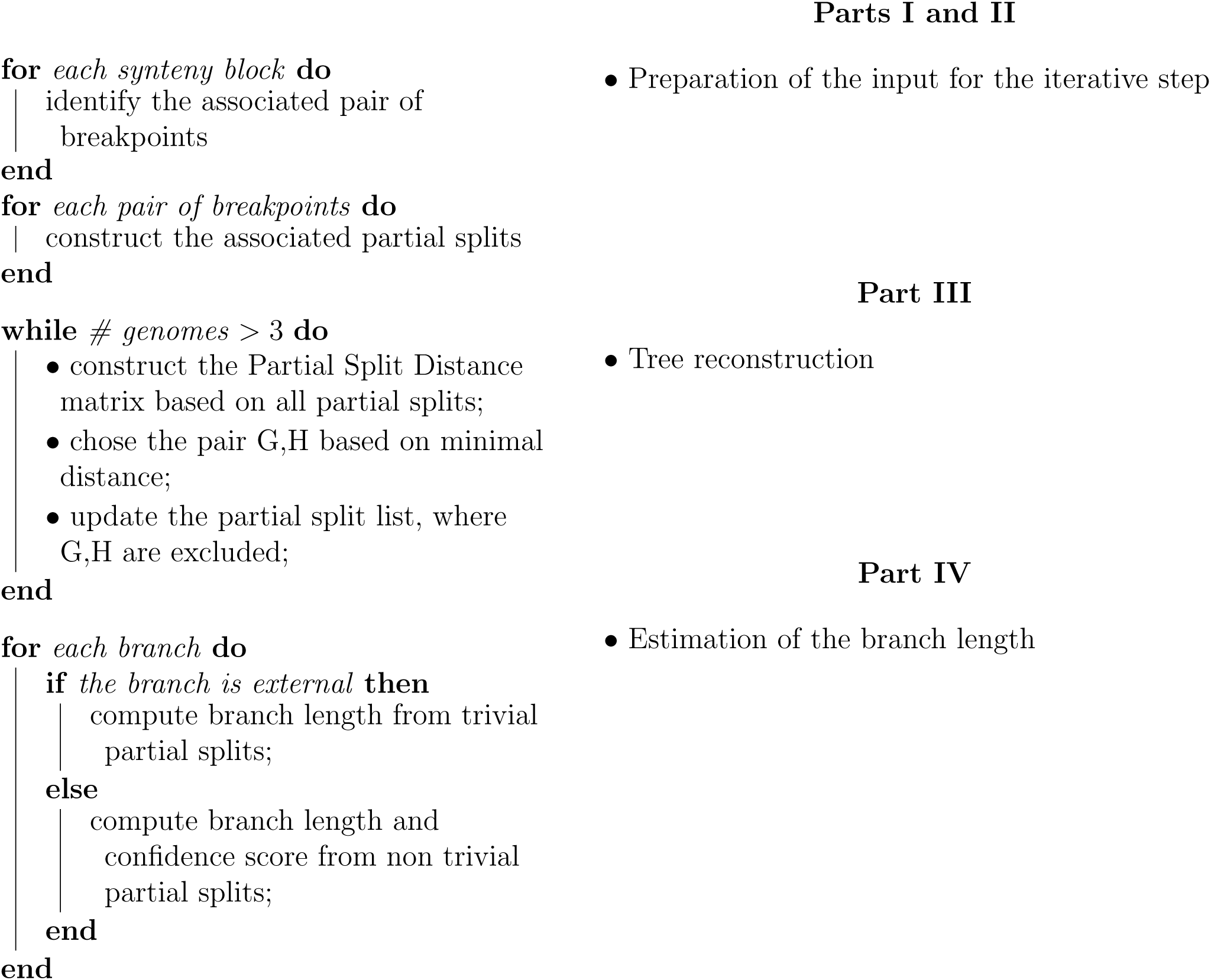
PhyChro algorithm.

#### Part I - Identification of breakpoints

*Step 1:* For each pairwise comparison *G/H* between pairs of genomes among *n* involved in the reconstruction, PhyChro iteratively identifies the breakpoints associated to each synteny block. See I in **Figure 4**.

#### Part II - Identification of partial splits

*Step 2:* For each breakpoint [(*B A*)_*G*_, (*B C*)_*H*_] identified in Step 1 and issued from the comparison *G/H* and for each genome *K* ≠ *G, H*, PhyChro determines whether (*B A*)_*G*_ or (*B C*)_*H*_ is present in *K* (as seen in section “Testing the conservation of block adjacencies”).

*Step 3:* Based on the results from Step 2, PhyChro defines two sets of genomes, 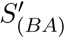 and 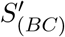, that share one or the other adjacency defining the breakpoint [(*B A*)_*G*_, (*B C*)_*H*_]. If 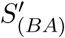 and 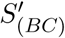 are not disjoint, then the sets are ignored (as seen in section “Partial split assigned to breakpoints”). These partial splits are associated to ambiguous breakpoints, which are themselves due to small blocks. If 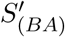 and 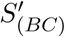 are disjoint, then PhyChro removes from the two sets those genomes that support only weakly the adjacency (as seen in section “Testing the conservation of block adjacencies”). Then it checks that both resulting sets *S*_(*BA*)_ and *S*_(*BC*)_ are not singletons; if so, it adds *S*_(*B A*)_‖*S*_(*B C*)_ to the collection of partial splits. Note that *S*_(*B A*)_‖*S*_(*B C*)_ may be trivial or not.

At the end of the iteration (steps 2 and 3), PhyChro has identified a collection of partial splits.

#### Part III - Bottom-up tree reconstruction

*Step 4:* For each pair of genomes *G, H*, PhyChro computes their PSD, f_inc_(*G, H*) and f_comp_(*G, H*) (as seen in section “The Partial Split Distance”).

*Step 5:* The creation of an internal node {*KL*} of the tree relies on the identification of the two sister genomes *K* and *L* (among the *n* genomes) displaying the smallest f_inc_ value. However, as explained above, to avoid considering very distant genomes that could have very small f_inc_ values, sister genomes are chosen to be the pair displaying the smallest f_inc_ value among the *n/*2 genome pairs that have the smallest ratio ℛ (III in **Figure 4**). Notice that the maximum number of possible sister genomes in a tree of *n* species is *n/*2. If there are multiple identical minimal f_inc_ values, either they involve different pairs of genomes and they will be treated one after the other in the different and successive iterations, or they involve incompatible pairs of genomes (involving the same genomes; a very unlikely situation that would results into the creation of a node with a low confidence score - see below) and the choice among them is left arbitrary.

*Step 6:* Once the internal node {*KL*} is created, the list of partial splits identified at Step 3, is updated by replacing all occurrences of *K* and *L* by the node {*KL*}. Two types of partial splits *S*_(*B A*)_‖*S*_(*B C*)_ are deleted: (i) partial splits that are discordant with the new node, that is partial splits where *K* and *L* belong to *S*_(*B A*)_ and *S*_(*B C*)_, respectively (see partial split 3 in III of **Figure 4**), (ii) partial splits characterized by a set of genomes composed by *K* and *L* only, since these partial splits would become trivial carrying no useful information for further topology reconstruction (see partial split 4 in III of **Figure 4**).

The process (steps 4-6) is iterated on the restricted set of genomes, where *K, L* are replaced by the ancestral genome {*KL*}, and on the updated set of partial splits obtained in step 6: all f_inc_ and f_comp_ values are re-computed from the updated list of partial splits, new internal nodes are created, and the list of partial splits is updated again. The iteration is run until only three genomes remain (exactly one unrooted tree topology is then possible).

#### Part IV. Estimations on the branches of the phylogenetic tree

PhyChro produces an estimation of the branch length and a confidence score of the reconstructed nodes. The branch length is an indicator of the complexity of the chromosomal structures (that is, of the amount of rearrangements identifiable from the genomes under consideration), and the confidence score indicates how much the reconstruction is supported and/or contradicted by the information contained in the initial non-trivial partial splits.

*Step 7:* Branch length for internal and terminal branches is estimated by using information contained in non-trivial and trivial partial splits, respectively. Branch length is the sum of a weighted number of partial splits (corresponding to a number of breakpoints, see **Supplementary File**) that support the existence of the branch (**Figure 6**), and therefore it indirectly represents a number of rearrangements. These values are necessarily an underestimation because most partial splits support the existence of a path in the tree rather than a specific branch, and therefore, are not considered for the calculation of branch lengths. In addition, terminal branches of distant genomes and internal branches between distant clades will be even more underestimated as partial splits supporting this kind of branches are rare.

**Figure 6.**
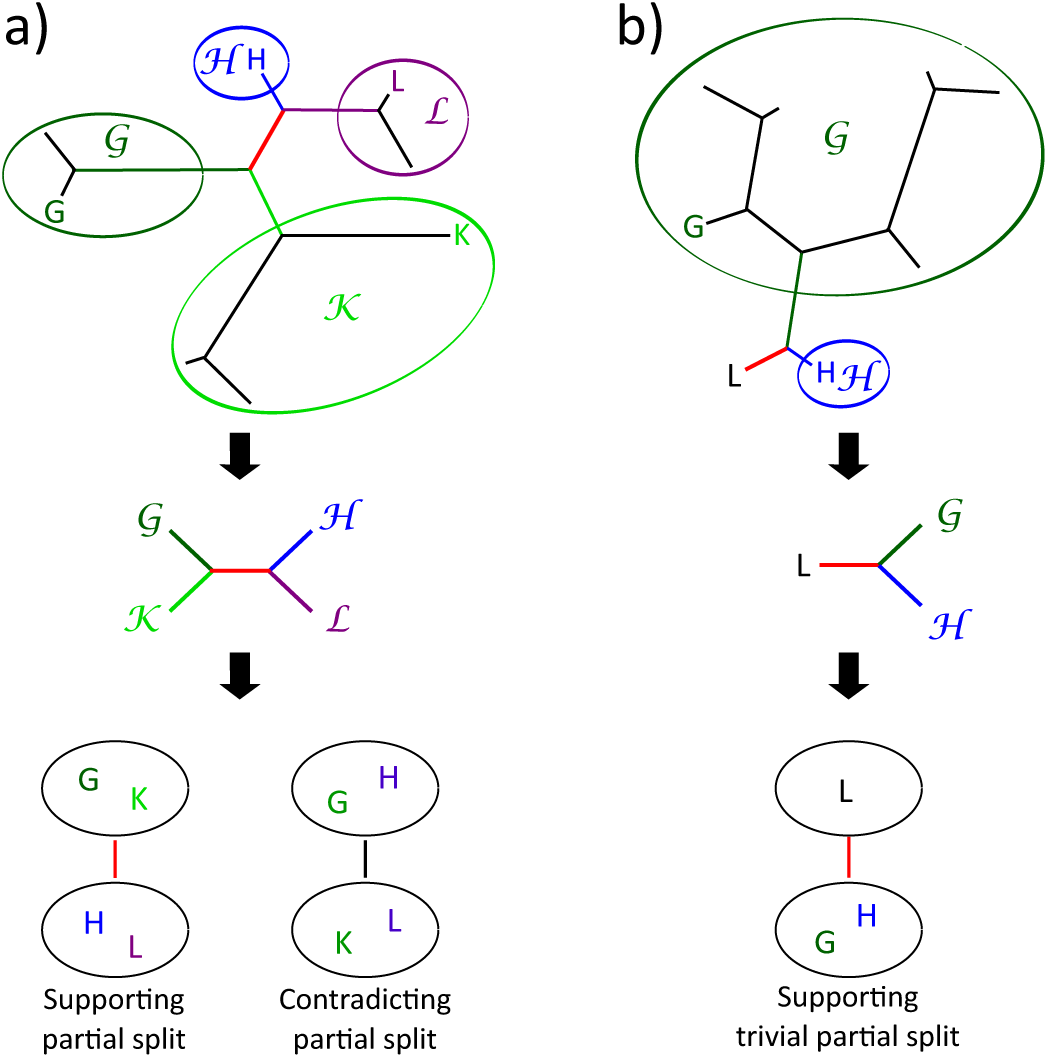
Examples of partial splits supporting or contradicting the existence of a given branch. Given a branch (red edges in the left and right trees), we consider the sets of genomes ℋ, 𝒢, 𝒦, ℒ corresponding to the maximal subtrees associated to the edge by the tree topology. Sets ℋ, 𝒢, 𝒦, ℒ contain genomes *H, G, K, L*, respectively. **a)** Each internal branch is characterized by a double pair of genome sets [(𝒢, 𝒦), (ℋ, ℒ)], which allows to define the partial splits that support or contradict this branch. **b)** Each external branch is characterized by one pair of genome sets (𝒢, ℋ), which allows to define the trivial partial splits that support this branch.

PhyChro also estimates a confidence score for each internal branch by calculating the proportion of non-trivial partial splits that supports its existence over the total number of non-trivial partial splits that either support or contradict it (**Figure 6** and **Supplementary File**). In addition to the confidence score, PhyChro provides the list of all *f*_*inc*_ values computed for genome pairs, which can help to know if a node is trustworthy or not.

### Description of input data

PhyChro requires as input the list of synteny blocks computed for each pairwise comparison *G/H* between all pairs of genomes *G, H* involved in the phylogenetic reconstruction. Anchors must be provided for each pair of synteny blocks issued from a comparison *G/H*. We recall that synteny blocks handled by PhyChro can overlap and that the same gene can be an anchor for distinct blocks. Duplicated synteny blocks are treated as independent blocks even though their anchors can be shared.

PhyChro accepts synteny blocks that are reconstructed with various tools as long as they are converted into the expected format, described in the README file of the PhyChro package. For the applications to yeast and vertebrate species, synteny blocks were computed with the SynChro software (Drillon et al., 2014), setting the Δ parameter to 3. Δ is a parameter that allows to define synteny blocks by controlling the complexity of internal micro-rearrangements. Intuitively, high values of Δ are more permissive and allow larger micro-rearrangements to be tolerated within synteny blocks while smaller values of Δ are more stringent and split synteny blocks at micro-rearrangement breakpoints. This implies that, for distantly related genomes, increasing the Δ value allows to recover a larger number of synteny blocks. For these genomes, small values of Δ would allow recovering the signal only from small inversions. Notice that when PhyChro reconstructs trees using blocks computed with Δ = 2, the yeast tree contains 3 erroneous splits and the vertebrate tree contains 1, while both trees are correct when blocks are computed with Δ = 3 or 4. SynChro automatically reconstructs pairwise synteny blocks that can be directly read by PhyChro, and it can be downloaded at www.lcqb.upmc.fr/CHROnicle/SynChro.html.

To analyze how sensitive is PhyChro to synteny block reconstruction, we constructed a second set of synteny blocks with the program i-ADHoRe 3.0 (Proost et al., 2012). We followed the protocol used in (Drillon et al., 2014) fixing parameters as follows: prob.cutoff=0.001, gap size=15, cluster.gap=20, q value=0.9 and anchor.points=3. The remaining parameters were set with default values. The i-ADHoRe 3.0 software package is available at bioinformatics.psb.ugent.be/software.

PhyChro was tested on 13 vertebrate species and 21 yeast species. The detailed list is given in the **Supplementary Table 1**. The vertebrate genome sequences have been downloaded from NCBI and the yeast species were downloaded from several sites listed in **Supplementary Table 2**.

### PhyChro computational time

PhyChro time complexity depends on the number of genomes given in input and on the number of rearrangements that took place among these genomes. Phylogenetic reconstructions with PhyChro ran, on a desk computer, in 15 and 10 minutes for the 13 vertebrate and 21 yeast species, respectively. A total of 130,485 and 179,649 breakpoints and of 17,848 (1,501 different ones) and 20,924 (3,901) partial splits were identified for vertebrate and yeast genomes, respectively.

### Comparison with MLGO

PhyChro has been compared with the method of phylogenetic reconstruction Maximum Likelihood for Gene Order Analysis (MLGO). MLGO’s input is constituted by chromosomes described as sequences of gene identifiers and these latter can be used multiple times, that is gene duplicates are allowed in MLGO. To prepare the input to MLGO, we used OrthoMCL as suggested in (Lin et al., 2013). Genes have been clustered using OrthoMCL (Li et al., 2003) with 1.5 as inflation value, 30% of similarity cut-off and a E-value of 10*e*-5. The same label has been used for genes falling in the same cluster. MLGO analysis was run at geneorder.com/server.php (Lin et al., 2013).

### Phylogenetic reconstructions based on protein sequences

We identified 357 families of syntenic homologs (considered as orthologs) sharing more than 90% of similarity between the 13 vertebrate species, and 80 families sharing more than 80% of similarity between the 21 yeast species, using SynChro (Drillon et al., 2014). Orthologous proteins were aligned with MUSCLE (version 3.8.31) (Edgar, 2004) and alignments were cleaned with Gblocks (version 0.91b) (Castresana, 2000). Cleaned concatenated alignments were then provided to PhyML 3.0 (which was run with the LG amino-acid substitution model) and ProtPars. For Neighbor, we computed the distance matrix using ProtDist and ran it with the neighbour joining option. ProtPars, Neighbor and ProtDist are included in the PHYLogeny Inference Package (version 3.67) (Felsenstein, 1989) and have been used online at mobyle.pasteur.fr/cgibin/portal.py.

## Acknowledgments

We thank Ingrid Lafontaine for critical reading of the manuscript. This work was supported by the Agence Nationale de la Recherche (‘GB-3G’, ANR-10-BLAN-1606-01) (GF), an ATIP-Avenir Plus grant from the CNRS (GF), a teaching assistantship (ATER) from the Ministère de l’Enseignement Supérieure et de la Recherche (GD), and funds from the Institut Universitaire de France (AC).

## Supplementary Material

### How to compute branch length and confidence score

#### For internal branches

Each internal branch *b* is flanked by four branches leading to four sets of genomes, two for each extremity of *b* (**Figure 6a**). This means that we can describe *b* by a double pair of genome sets [(𝒢, 𝒦), (ℋ, 𝒬)], where 𝒢 and 𝒦 are two sets of genomes lying in the maximal subtree excluding *b* and containing one of its extremities, and where ℋ and 𝒬 are the two sets of genomes lying in the maximal subtree excluding *b* and containing the other extremity of *b*. Depending on these two pairs of sets, we can determine whether a given partial split *S*_(*B A*)_ ‖*S*_(*B C*)_ supports the branch *b* or not. To do so, we define the predicate p_support_ as follows:

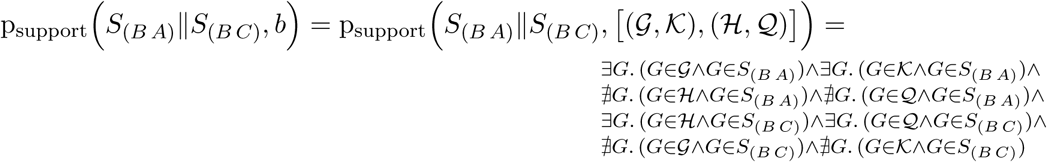

and say that *S*_(*B A*)_‖*S*_(*B C*)_ supports *b* if the predicate is true. The left panel of **Figure 6a** gives an example of a partial split that supports a given branch.

Similarly, we can determine when a given partial split contradicts a given internal branch *b*, meaning that this partial split would imply at least two rearrangements occurring in two different branches of the tree and involving the same block extremities. To do so, we define the predicate p_contradict_ as follows:

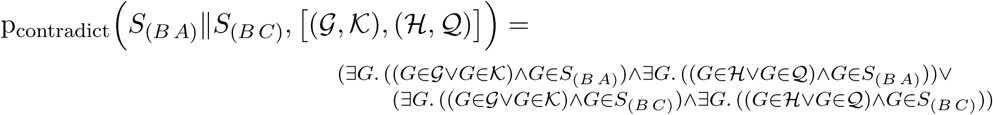

and say that *S*_(*B A*)_‖*S*_(*B C*)_ contradicts *b* if the predicate is true. The right panel of **Figure 6a** gives an example of a partial split that contradicts a given branch.

For each internal branch *b*, we define the confidence score of *b* as:

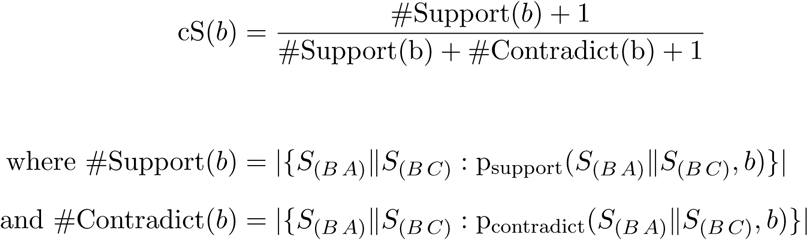

Also, we would like to compute the length of an internal branch as the number of breakpoints that would be identified if the two ancestral genomes associated to the extremities of *b* were compared. To estimate this number, we use the predicate p_support_, since every partial split supporting *b* is indeed associated to one of these breakpoints. However, the calculation is not that simple, since each breakpoint is associated to several partial splits rather than to a single one. For instance, given two sets of genomes, *S*_(*B A*)_ and *S*_(*B C*)_, regrouping the actual genomes that have conserved the two adjacencies of a given breakpoint [(*B A*)_*G*_, (*B C*)_*H*_] respectively, each comparison between two genomes from these two sets should lead to the identification of a partial split associated to this breakpoint. There are |*S*_(*B A*)_| * |*S*_(*B C*)_| such comparisons. Therefore, the number of breakpoints is estimated from the number of partial splits by weighting them according to the maximal number of partial splits that could be associated to the same breakpoint:

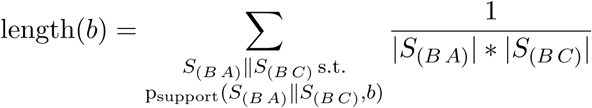

#### For terminal branches

Each terminal branch, leading to a genome *K*, is characterized by two branches leading to two sets of genomes (**Figure 6b**), and therefore, it can be described by a pair [*K*, (𝒢, ℋ)], where 𝒢 and ℋ are the two sets of genomes. Depending on these sets, we can determine whether a given trivial partial split, *S*_(*B A*)_‖*S*_(*B C*)_, where *S*_(*B A*)_ is constituted by an unique genome, supports this branch or not. This is done with the predicate t_support_ defined as follows:

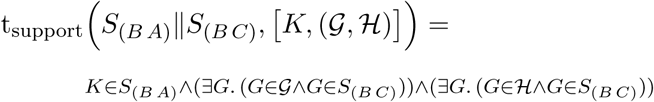

We say that *S*_(*B A*)_‖*S*_(*B C*)_ supports the external branch *b* if the predicate is true. **Figure 6b** gives an example of trivial partial split that supports a given external branch.

From this predicate, we compute the length of an external branch *b* as follows:

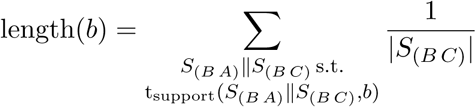

## Supplementary Tables

**Table S1:** List of 21 yeasts and 16 vertebrates with a high quality assembled genome.

| Class | Species | Genome size<br>(Mb) | # of<br>Chr. | # of<br>Scaf. | # of<br>Gen. | Reference |
| --- | --- | --- | --- | --- | --- | --- |
| Saccharomycetes | *Candida albicans* | 14.3 | 8 | 8 | 6182 | (Jones *et al.*, 2004) |
| Saccharomycetes | *Candida dubliniensis* | 14.6 | 8<sup>1</sup> | 8<sup>1</sup> | 5858 | (Jackson *et al.*, 2009) |
| Saccharomycetes | *Candida glabrata* | 12.3 | 13 | 13 | 5202 | (Dujon *et al.*, 2004) |
| Saccharomycetes | *Candida parapsilosis* | 13.1 | 7 | 14 | 5608 | (Butler *et al.*, 2009) |
| Saccharomycetes | *Candida tropicalis* | 14.6 | 8 | 20 | 6253 | (Butler *et al.*, 2009) |
| Saccharomycetes | *Clavispora lusitanae* | 12.1 | 8 | 8 | 5936 | (Butler *et al.*, 2009) |
| Saccharomycetes | *Debaryomyces hansenii* | 12.2 | 7 | 7 | 6272 | (Dujon *et al.*, 2004) |
| Saccharomycetes | *Eremothecium gossypii* | 8.7 | 7 | 7 | 4768 | (Dietrich *et al.*, 2004) |
| Saccharomycetes | *Kluyveromyces lactis* | 10.7 | 6 | 6 | 5076 | (Dujon *et al.*, 2004) |
| Saccharomycetes | *Lachancea kluyveri* | 11.3 | 8 | 8 | 5321 | (Souciet *et al.*, 2009) |
| Saccharomycetes | *Lachancea thermotolerans* | 10.4 | 8 | 8 | 5092 | (Souciet *et al.*, 2009) |
| Saccharomycetes | *Lachancea waltii* | 10.7 | 8 | 10 | 6614 | (Kellis *et al.*, 2004) |
| Saccharomycetes | *Lodderomyces elongisporus* | 15.5 | 8 | 22 | 5795 | (Butler *et al.*, 2009) |
| Saccharomycetes | *Naumovozyma castellii* | 11.2 | 9 | 10 | 5648 | (Gordon *et al.*, 2011) |
| Saccharomycetes | *Pichia guilliermondii* | 10.6 | 8 | 9 | 5920 | (Butler *et al.*, 2009) |
| Saccharomycetes | *Pichia pastoris* | 9.4 | 4 | 6 | 5077 | (De Schutter *et al.*, 2009) |
| Saccharomycetes | *Pichia stipitis* | 15.4 | 8 | 9 | 5818 | (Jeffries *et al.*, 2007) |
| Saccharomycetes | *Saccharomyces cerevisiae* | 12.1 | 16 | 16 | 6664 | (Goffeau *et al.*, 1996) |
| Saccharomycetes | *Torulaspora delbrueckii* | 9.2 | 6 | 8 | 4972 | (Gomez-Angulo *et al.*, 2015) |
| Saccharomycetes | *Yarrowia lipolytica* | 20.5 | 6 | 6 | 6448 | (Dujon *et al.*, 2004) |
| Saccharomycetes | *Zygosaccharomyces rouzii* | 9.8 | 7 | 7 | 4991 | (Souciet *et al.*, 2009) |
| Sauropsida | *Anolis carolinensis* | 1780 | 18 | 18 | 18595 | (Alfoldi *et al.*, 2011) |
| Mammalia | *Bos taurus* | 2500 | 30 | 30 | 24293 | (Zimin *et al.*, 2009) |
| Mammalia | *Canis familiaris* | 2400 | 39 | 39 | 19014 | (Lindblad-Toh *et al.*, 2005) |
| Actinopterygii | *Danio rerio* | 1700 | 25 | 25 | 22940 | (Howe *et al.*, 2013) |
| Mammalia | *Equus caballus* | 2689 | 32 | 32 | 20257 | (Wade *et al.*, 2009) |
| Aves | *Gallus gallus* | 1000 | 40<sup>2</sup> | 29 | 15308 | (The International Chicken Genome Sequencing Consortium, 2004) |
| Mammalia | *Homo sapiens* | 3080 | 23 | 23 | 19439 | (The International Human Genome Sequencing Consortium, 2001) |
| Mammalia | *Macaca mulatta* | 2871 | 22 | 21 | 21023 | (The Rhesus Macaque Genome Sequencing and Analysis Consortium, 2007) |
| Marsupialia | *Monodelphis domestica* | 3475 | 9 | 9 | 18640 | (Mikkelsen *et al.*, 2007) |
| Mammalia | *Mus musculus* | 2644 | 20 | 20 | 21923 | (The Mouse Genome Sequencing Consortium, 2002) |
| Actinopterygii | *Oryzias latipes* | 800 | 24 | 24 | 17445 | (Kasahara *et al.*, 2007) |
| Mammalia | *Pan troglodytes* | 3100 | 24 | 24 | 19125 | (The Chimpanzee Sequencing and Analysis Consortium, 2005) |
| Mammalia | *Rattus Norvegicus* | 3000 | 21 | 21 | 22925 | (The Rat Genome Sequencing Project Consortium, 2004) |
| Mammalia | *Sus scrofa* | 2800 | 19 | 19 | 21630 | (Wernersson *et al.*, 2005) |
| Aves | *Taeniopygia guttata* | 2644 | 28 | 28 | 12337 | (Warren *et al.*, 2010) |
| Actinopterygii | *Tetraodon nigroviridis* | 350 | 21 | 21 | 13580 | (Jaillon *et al.*, 2004) |

<sup>a</sup>Pseudochromosomes obtained by mapping onto *C. albicans* chromosomes (Jackson *et al.*, 2009).<sup>b</sup>Including microchromosomes that were not assembled.

**Table S2:**
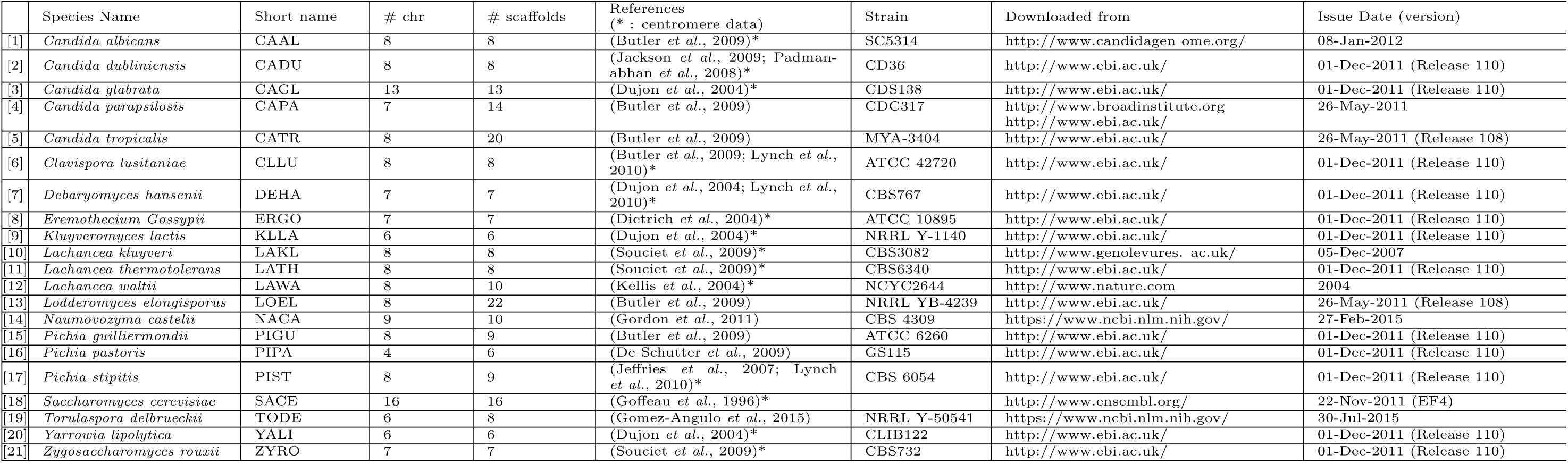
21 Yeast Species Summary

## Supplementary Figures

**Figure S1:**
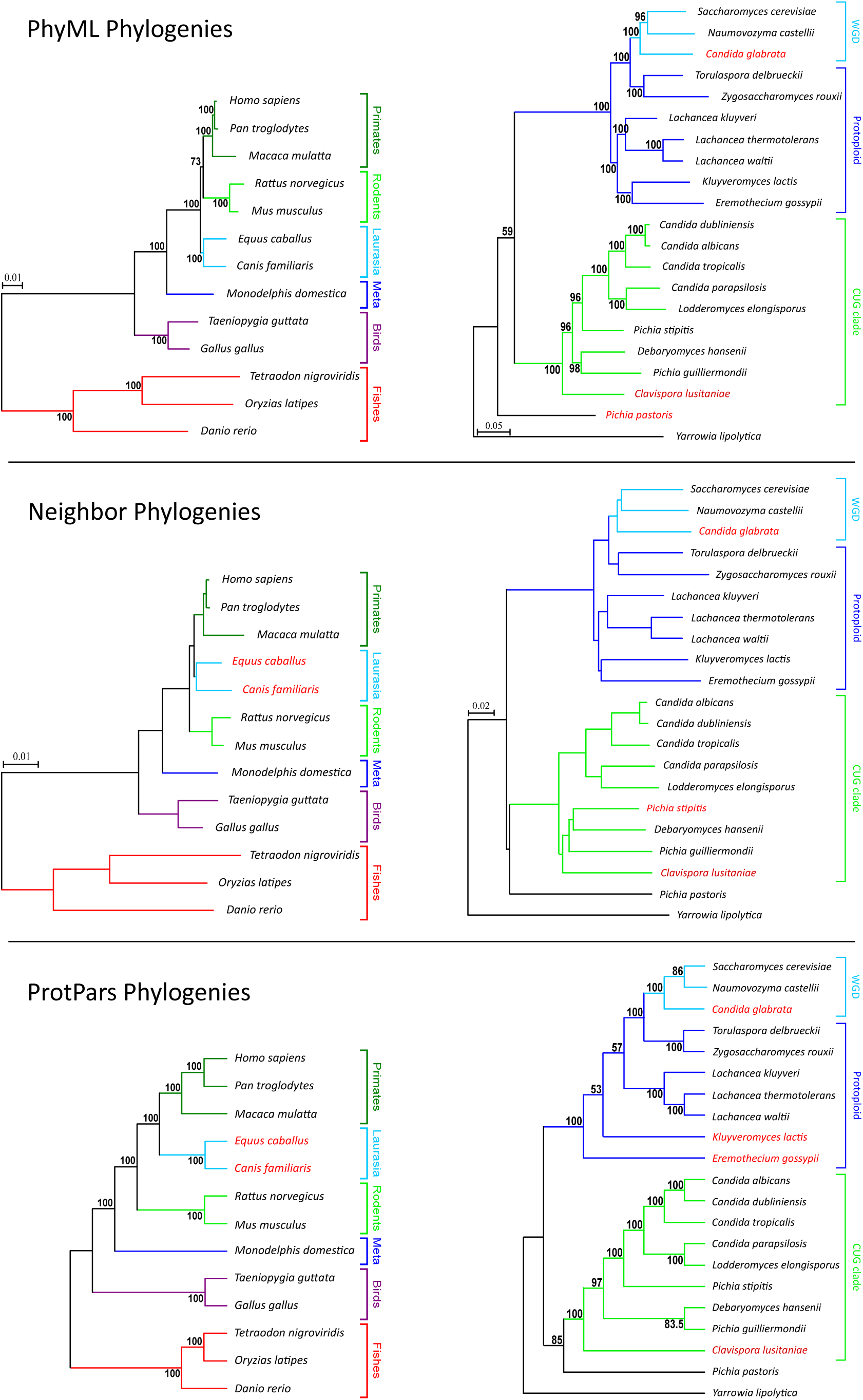
Phylogenies obtained with PhyML, Neighbor and ProtPars methods. Bootstrap values are given for the PhyML and the ProtPars phylogenies (for 100 resampled data sets). For PhyML and Neighbor reconstructions, branch length scales are indicated on the left of the trees.

**Figure S2:**
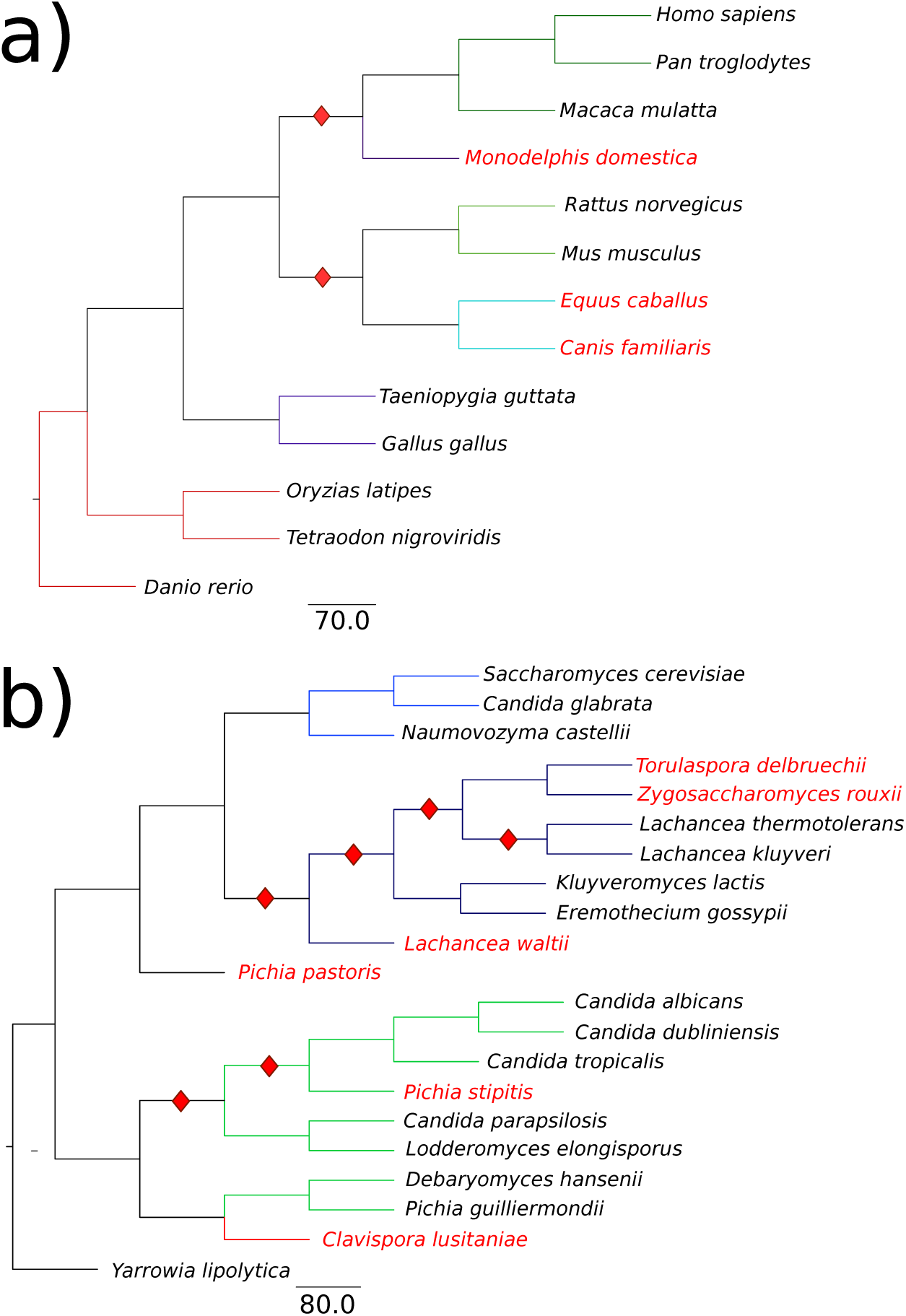
Phylogenies obtained with MLGO method. Tree reconstructions realized with MLGO, for vertebrates (**a**) and yeasts (**b**). Red losanges indicate erroneous splitting of the tree compared to the known one. Compare to **Figure 2**, where the colors assigned to branches are the same.

**Figure S3:**
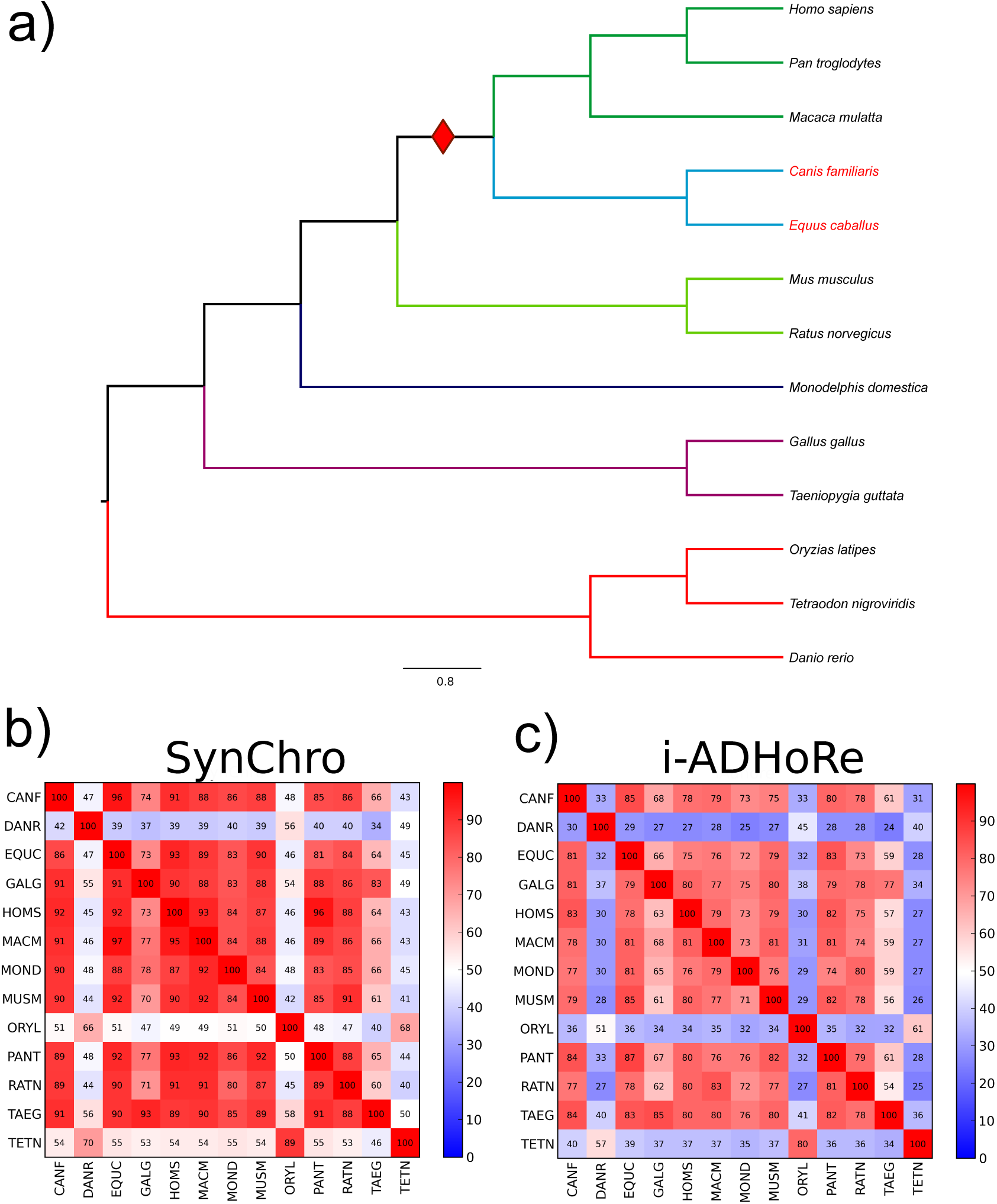
Vertebrate phylogeny obtained with PhyChro based on i-ADHoRe synteny blocks. **a)** Tree topology reconstruction realized with PhyChro and based on i-ADHoRe synteny blocks. The red losange indicates an error in the phylogenetic reconstruction. Colors correspond to the ones used in **Figure 2a**. Compare it to **Figure 2a. b)** Matrix representing the synteny blocks coverage of the vertebrate genomes after pairwise comparison, where synteny blocks are obtained with SynChro (run with Δ = 3). At row X and column Y, the number in the cell of the matrix corresponds to the coverage of genome Y after comparison with genome X, that is the percentage of the number of genes in the genome that belong to synteny blocks. The scale indicates the coverage level and goes from low (blue) to high (red). **c)** Matrix representing the synteny blocks coverage of the vertebrate genomes after pairwise comparison, where synteny blocks are obtained with i-ADHoRe.

**Figure S4:**
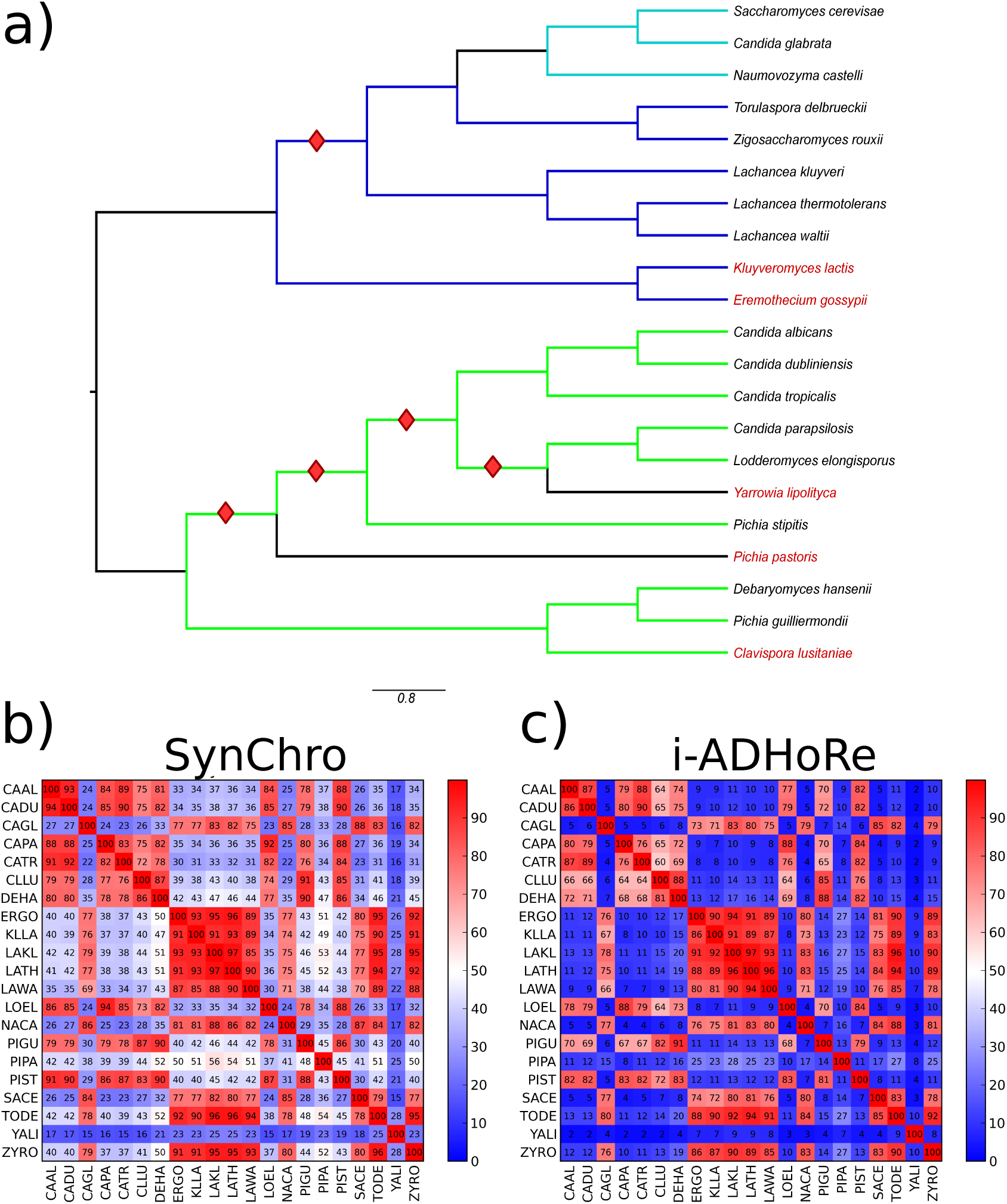
Yeasts phylogeny obtained with PhyChro based on i-ADHoRe synteny blocks. **a)** Tree topology reconstruction realized with PhyChro and based on i-ADHoRe synteny blocks. Red losanges indicate errors in the phylogenetic reconstruction. Colors correspond to the ones used in **Figure 2b**. Compare it to **Figure 2b. b)** Matrix representing the synteny blocks coverage of the yeast genomes after pairwise comparison, where synteny blocks are obtained with SynChro (run with Δ = 3). At row X and column Y, the number in the cell of the matrix corresponds to the coverage of genome Y after comparison with genome X, that is the percentage of the number of genes in the genome that belong to synteny blocks. The scale indicates the coverage level and goes from low (blue) to high (red). **c)** Matrix representing the synteny blocks coverage of the yeast genomes after pairwise comparison, where synteny blocks are obtained with i-ADHoRe.

**Figure S5:**
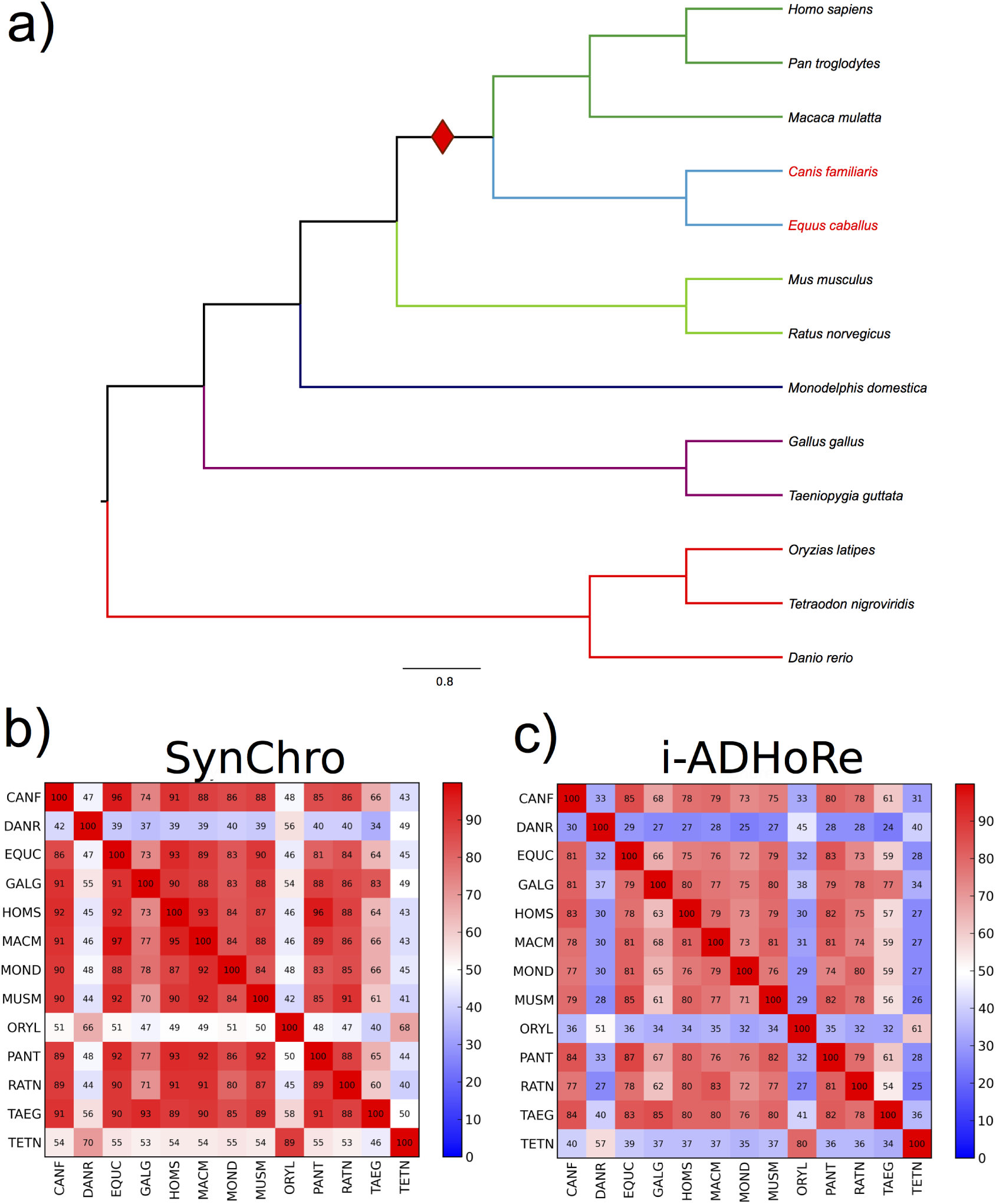
Distribution of blocks obtained by i-ADHoRe and SynChro on vertebrate and yeast species. The *x*-axis reports the number of genes in a block (block size) and the *y*-axis reports the number of blocks of a given size found in all pairwise comparisons between vertebrate (left) and yeast (right) species. Blocks of size ≥ 21 are added up in the last columns for both i-ADHoRe (red) and SynChro (black).

**Figure S6:**
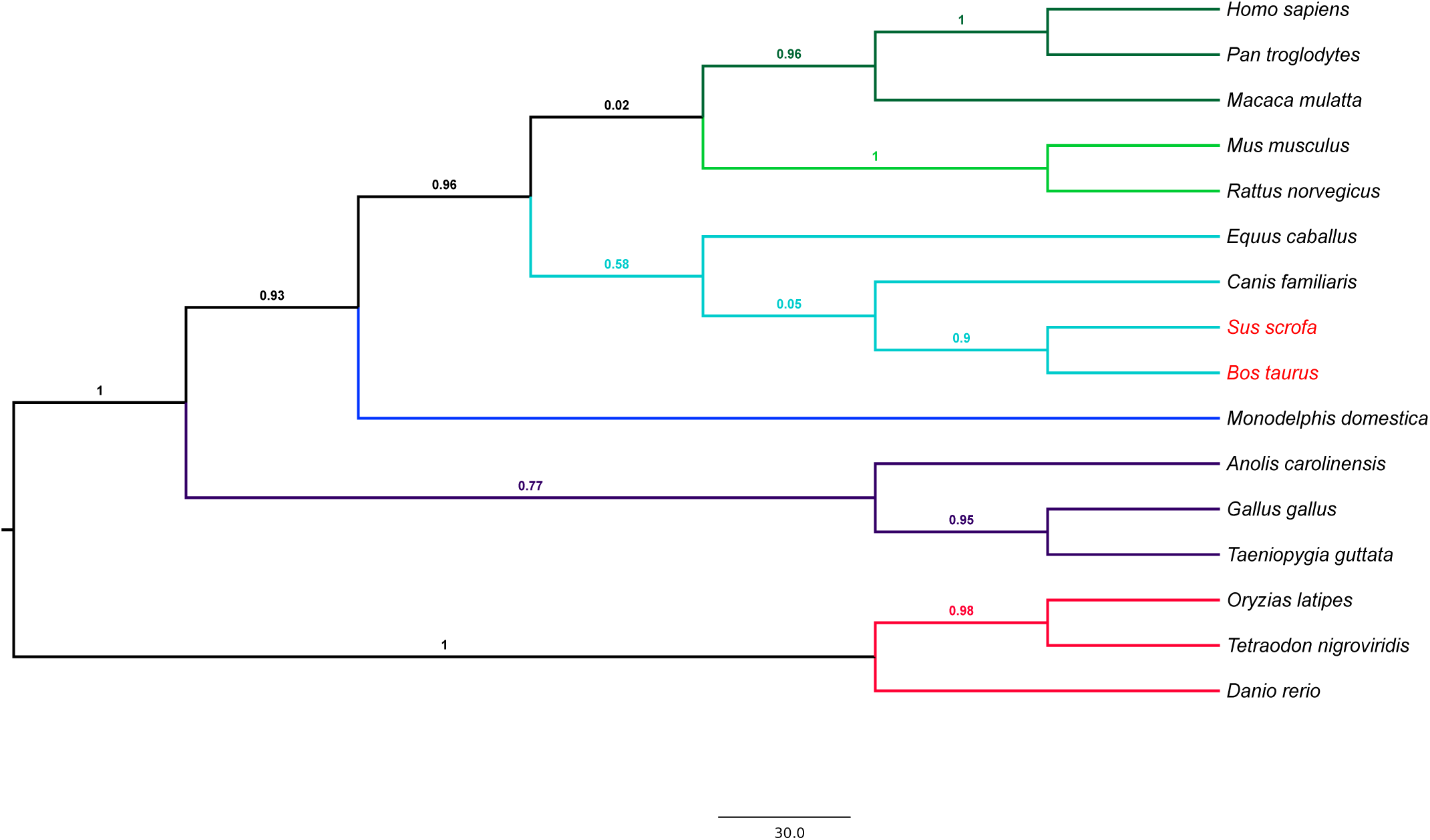
Phylogenies obtained with PhyChro for the 13 vertebrates plus the cow, the pig and the lizard. See legend in **Figure 2**. In this reconstruction, the position of horse and dog is not well supported by PhyChro confidence score (0.05) and this suggests the possibility of alternative topological nestings. By looking at the data produced by PhyChro partial split analysis for the reconstruction of the tree in **Figure 2a**, note that horse and dog are sister genomes with *f*_*inc*_(dog,horse)= 38, a value that is much larger than what is ideally expected for sister genomes, that is 0. It corresponds to a large number of partial splits separating them, suggesting that the dog and the horse might have been particularly sensitive to convergent rearrangements (homoplasy) or to the accumulation of small inversions. Therefore, even if we assume that they actually are sister genomes in the tree including the cow and the pig, they are likely to be found further away from each other than they are from the cow and the pig. Indeed, this is what PhyChro finds: *f*_*inc*_(dog,horse)= 38, *f*_*inc*_((cow,pig),dog)= 10 and *f*_*inc*_((cow,pig),horse)= 21. Note that in (Romiguier et al., *Molecular Biology and Evolution*, 2013), the position of ((cow,pig),(dog,horse)) is well supported. This case illustrates well, on the one hand, the limits of PhyChro, which is sensitive to rearrangement convergences when such events exist (note that from the very good quality of PhyChro tree reconstructions, it appears that such rearrangement convergences are few), and on the other hand, its power, since its output can help to build a good understanding of the tree topology, and most of all, whether one should have confidence or not in the reconstructed topology.

**Figure S7:**
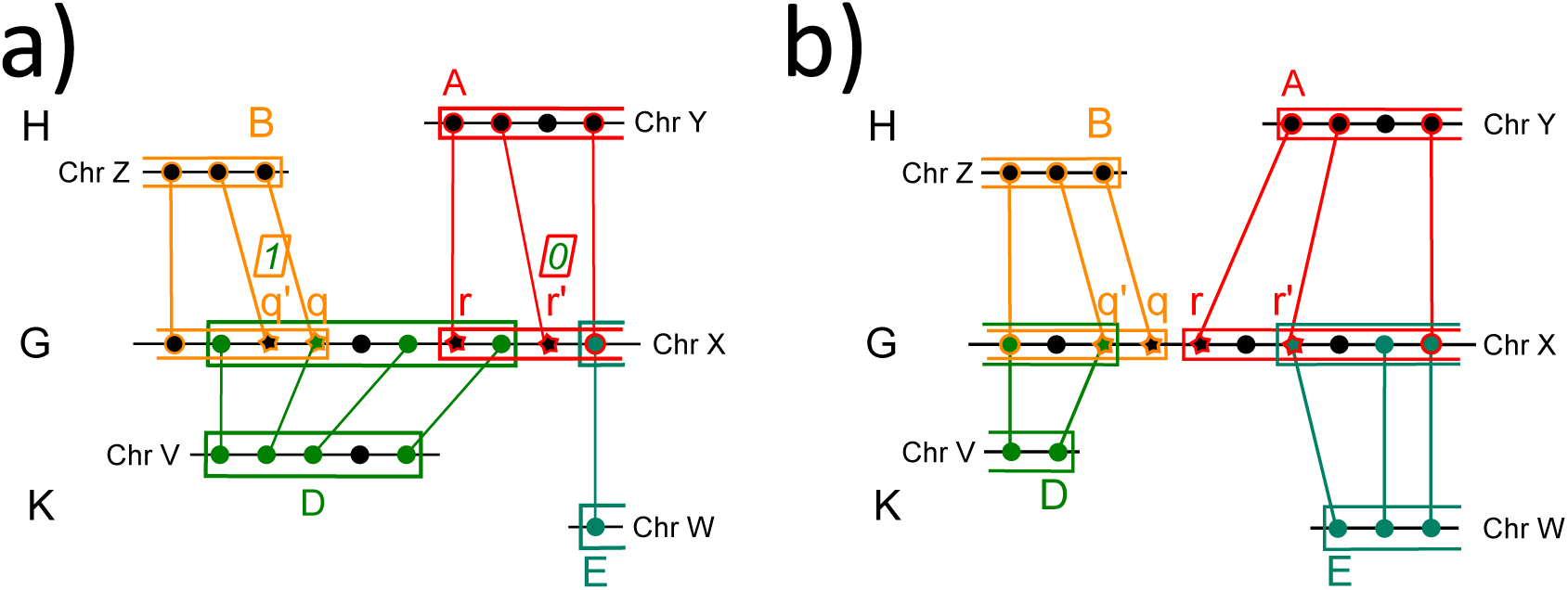
Illustration of two possible localizations of an adjacency (*B A*)_*G*_ in a genome *K*. Genes are indicated by dots or stars. Stars, in *G*, are used for the two first (*q*′ and *q*) and last (*r* and *r*′) anchors of blocks *B* and *A* respectively, in the comparison *G/H*. Red, yellow and green colors are used to highlight anchors associated to the blocks *A, B* and *D*, obtained in the comparisons *G/H, G/H* and *G/K*, respectively. **a)** (*BA*)_*G*_ ∈ *K* but *K* supports only weakly (*BA*)_*G*_: genes *q* and *r* belong to the same block *D*, along *G*, in *G/K* but the list of expected conditions is not completely fulfilled (see text). The number of anchors of *D* lying before *q*′ (after *r*′), and possibly including it, is indicated above *q*′ (*r*′) within a square. **b)** (*BA*)_*G*_ ∉ *K*: genes *q* and *r* do not belong to the same block in *G/K*.

**Figure S8:**
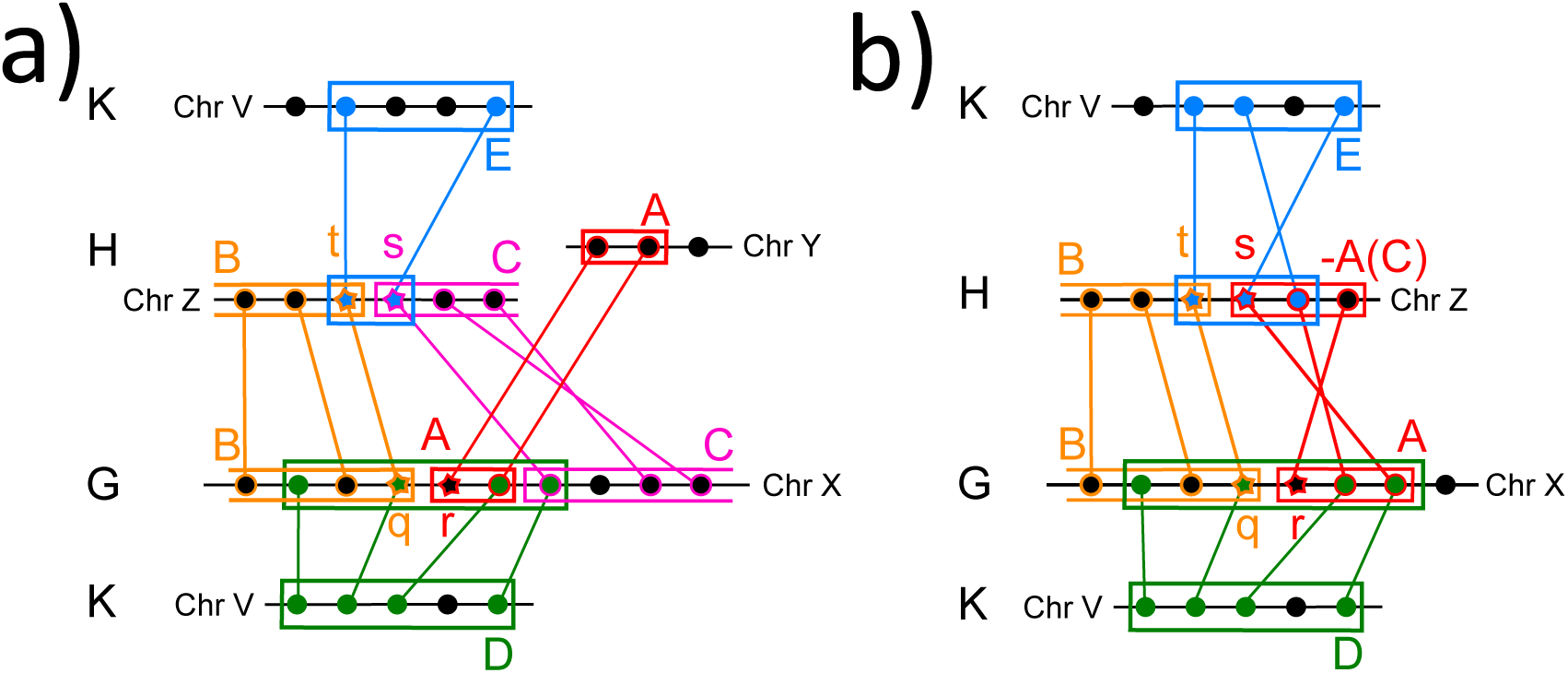
Illustration of two cases where *K* belongs to both sets *S*_*(B A)*_ and *S*_*(B C)*_. For sake of clarity, the 5 genes of chromosome *V* in genome *K*, which are involved in the adjacencies (*B A*)_*G*_ and (*B C*)_*H*_, are represented twice to illustrate both the *G/K* comparison (green) and the *H/K* comparison (blue). In the *G/K* (resp. *H/K*) comparison, genes *q, r* (resp. *t, s*) characterizing (*B A*)_*G*_ (resp. (*B C*)_*H*_), are included in the same block *D* (resp. *E*) along *G* (resp. *H*). **a)** Illustration of the ambiguous breakpoint [(*B A*)_*G*_, (*B C*)_*H*_], where *C* follows *A* in *G* and where *A* is a small block. **b)** Illustration of the ambiguous breakpoint [(*B A*)_*G*_, (*B* − *A*)_*H*_], where *A* is a small block.

